# Deletion of endocannabinoid synthesizing enzyme DAGLα from cerebellar Purkinje cells decreases social preference and elevates anxiety

**DOI:** 10.1101/2024.08.08.607068

**Authors:** Gabriella Smith, Kathleen McCoy, Gonzalo Viana Di Prisco, Alexander Kuklish, Emma Grant, Mayil Bhat, Sachin Patel, Ken Mackie, Brady Atwood, Anna Kalinovsky

## Abstract

The endocannabinoid (eCB) signaling system is robustly expressed in the cerebellum starting from the embryonic developmental stages to adulthood. There it plays a key role in regulating cerebellar synaptic plasticity and excitability, suggesting that impaired eCB signaling will lead to deficits in cerebellar adjustments of ongoing behaviors and cerebellar learning. Indeed, human mutations in *DAGLα* are associated with neurodevelopmental disorders. In this study, we show that selective deletion of the eCB synthesizing enzyme diacylglycerol lipase alpha (Daglα) from mouse cerebellar Purkinje cells (PCs) alters motor and social behaviors, disrupts short-term synaptic plasticity in both excitatory and inhibitory synapses, and reduces Purkinje cell activity during social exploration. Our results provide the first evidence for cerebellar-specific eCB regulation of social behaviors and implicate eCB regulation of synaptic plasticity and PC activity as the neural substrates contributing to these deficits.

Graphical abstract.
Cerebellar anatomy, morphology of Purkinje cells, localization, density, and spontaneous activity of excitatory and inhibitory synapses are normal in cerebellar-Purkinje-cell-specific Daglα KOs. However, endocannabinoid-dependent short-term synaptic plasticity (DSE and DSI) and activity of Purkinje cells in lobe VI during social exploration are dramatically reduced, and the KO mice exhibit alterations in sensorimotor coordination, deceased social preference, and increased anxiety.

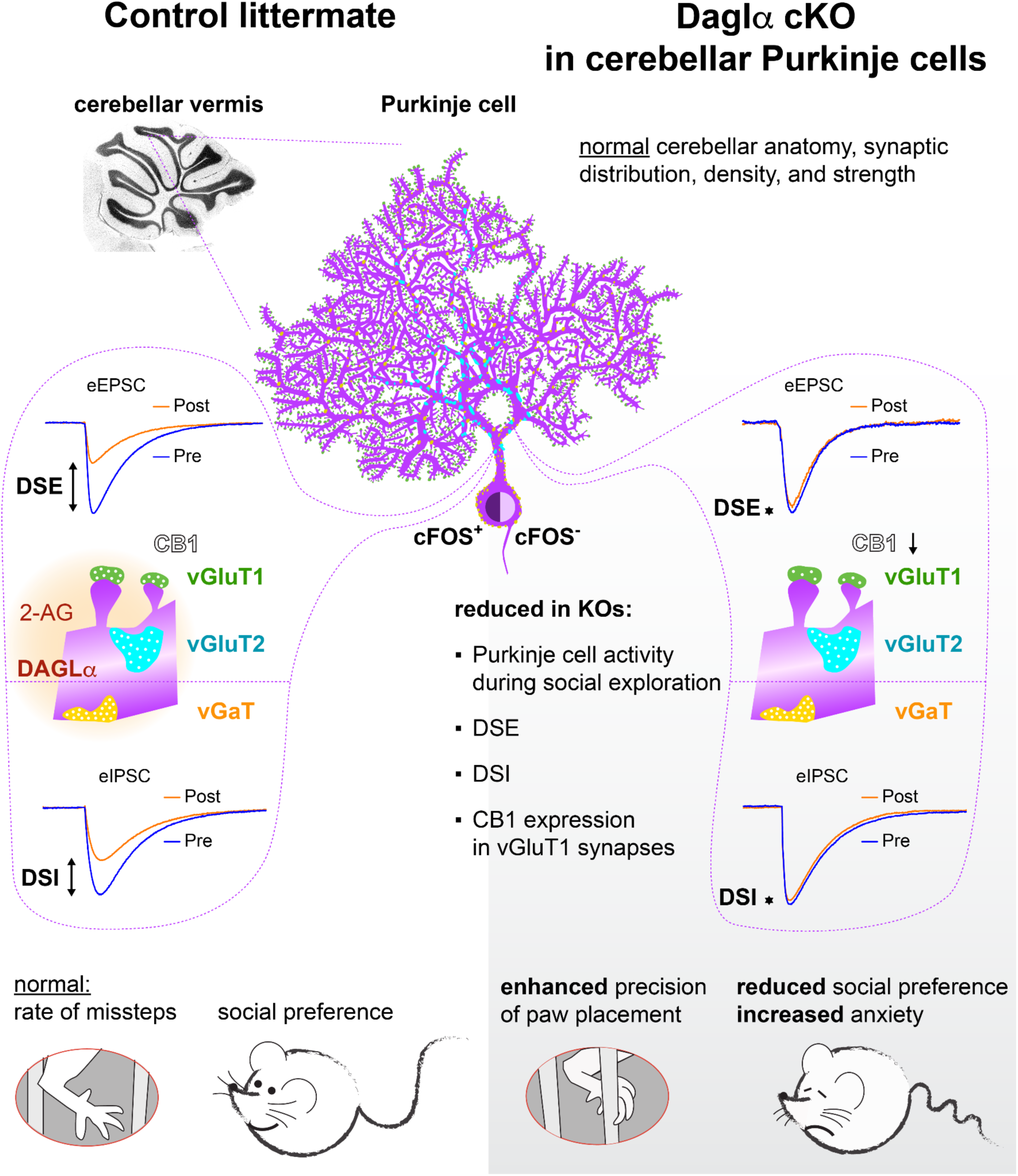

## INTRODUCTION

The cerebellum plays a key role in the regulation of implicit behaviors, ranging from motor coordination and balance to emotional processing [1][2][3][4]. Cerebellar outputs are controlled by Purkinje cell (PC) activity, which, in turn, is influenced by endocannabinoid (eCB) signaling [5][6][7][8]. eCB signaling machinery is prominently expressed in the cerebellum: diacylglycerol lipase alpha (Daglα), the primary synthesizing enzyme of 2-Arachidonoylglycerol (2-AG), a major neuronal eCB, is highly expressed in cerebellar PCs [9,10]; and cannabinoid receptor 1 (CB1), the main neuronal receptor for 2-AG, is expressed in the axons and the presynaptic terminals of both excitatory and inhibitory synaptic inputs to PCs [11–14].

Synaptic plasticity – changes in synaptic strength that persist on a wide range of timescales, from milliseconds-minutes (short-term) to hours-days-years (long-term) – is the main substrate of learning and memory [15]. eCB signaling plays a key role in cerebellar long-term and short-term synaptic plasticity, referred to as long-term synaptic potentiation and depression (LTP and LTD) [16,17] and depolarization-induced suppression of excitation and inhibition (DSE and DSI), respectively [6,11,12]. It is tempting to suggest that eCB signaling could be a key mediator of cerebellar-dependent learning and behavior.

Mutations in *DAGLα* are associated with neurodevelopmental disorders and cerebellar ataxias in humans [18][19,20] and are implicated in one of the most common neurodevelopmental diagnoses, Autism Spectrum Disorder (ASD) [21–24]. The criteria for ASD diagnosis [25] include reduced social interactions, increased repetitiveness, and atypical sensorimotor learning and performance – such as toe walking in toddlers, increased movement stereotypy, and sometimes extraordinary performance of technical and cognitive skills [26,27].

Cerebellar involvement in ASD pathology is supported by multiple lines of evidence. Developmental cerebellar lesions dramatically increase the predisposition for ASD [28] and PC loss is a common finding in ASD patients post-mortem [29]. PC-specific knockout mouse models of ASD-associated widely-expressed genes, such as FMR1, SHANK2, and TSC1/2, exhibit reduced sociability and other ASD-associated behavioral phenotypes [30–33], suggesting a critical role for the cerebellum in mediating pathology associated with their mutations. However, the role of cerebellar-PC-derived 2-AG in the regulation of social behaviors is unknown. To address this question, we generated PC-specific Daglα conditional knockout (KO) mice to explore the function of Daglα in cerebellar PC synaptic development and plasticity and in cerebellar-influenced behaviors.

Our results show alterations in motor adjustments during horizontal ladder locomotion, social approach, and anxiety in the KO mice, while forelimb coordination and repetitive behaviors are unaffected, implicating cerebellar eCB signaling in the regulation of a specific subset of behavioral domains. Confocal microscopy and 3D volumetric reconstruction were used to evaluate synaptic distribution in Daglα KO PCs, and electrophysiology in cerebellar slices was used to evaluate short-term synaptic plasticity. Daglα KO PCs exhibit reduced CB1 expression in vGluT1-positive presynaptic terminals, while the overall distribution and density of presynaptic terminals onto PC dendrites and somata are normal. Cannabinoid-dependent short-term synaptic plasticity in both excitatory and inhibitory synapses is dramatically reduced in Daglα KO PCs. The overall activity of Daglα KO PCs during social exploration, as assessed by expression of the immediate early gene cFos, is reduced concomitantly with reduced preference to explore social cues.

## RESULTS

### Daglα is selectively deleted from postnatal cerebellar Purkinje cells utilizing a Purkinje cell specific Cre mouse line

We generated a Purkinje cell specific Daglα KO mouse line by crossing the conditional floxed *Daglα* (*Daglα^fl^*) mouse generated by the group of Dr. Sachin Patel [34] with Purkinje cell specific Cre mouse (*Pcp2^Cre^*) generated by the group of Noboru Suzuki [35]. Littermate controls (LM, genotype *Pcp2^Cre/Cre^; Ai9^f/f^; Daglα^WT/WT^*) and conditional KOs (genotype *Pcp2^Cre/Cre^; Ai9^f/f^; Daglα^fl/fl^*) were generated by breeding males and females heterozygous for *Daglα^fl^* alle (*Pcp2^Cre/Cre^; Ai9^f/f^; Daglα^fl/WT^)* with tdTomato (TOM, Ai9) reporter mice (The Jackson Laboratory, https://www.jax.org/strain/007909) to allow for the visualization of recombined Purkinje cells via TOM expression (Fig. 1 A). Both female and male mice were used in all experiments reported below, but sex-combined results are shown for the experiments where we observed no sex differences.

**Figure 1.**
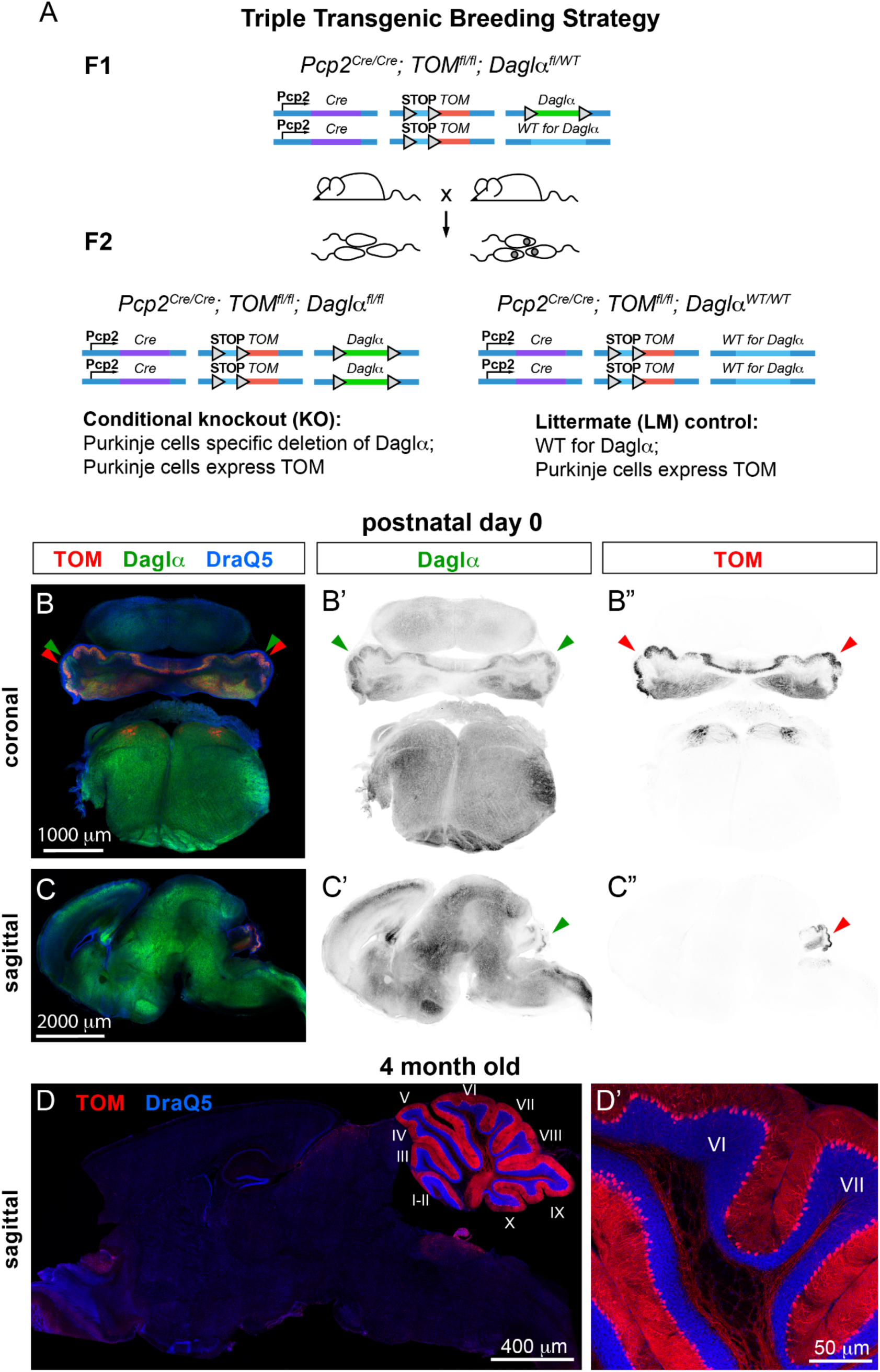
Pcp2^Cre^ recombination is efficient at birth and selective to Purkinje cells. (A) Littermate controls (LM), and conditional knockouts (KO) were generated by breeding Pcp2^Cre/Cre^; Ai9^fl/fl^; DAGLα^fl/WT^. The floxed tdTomato (TOM) allele (Ai9 reporter) was included, allowing the visualization of recombined Purkinje cells with TOM expression. Triangles represent LoxP sites. F1 = first generation, breeding; F2 = offspring, the genotypes used in experiments. (B-B’’) Coronal and (C-C”) midsagittal sections from newborn (postnatal day 0, P0) mice. The green arrows highlight the prominent expression of Daglα in the cerebellar Purkinje cells. Red arrows point to TOM reporter expression, verifying that Cre recombination is efficient and specific in the cerebellar Purkinje cells at birth. (D) The midsagittal section from a two-month-old mouse and (D’) higher magnification of cerebellar lobules VI-VII show that reporter expression is highly restricted to the cerebellar Purkinje cells.

At birth, Daglα is robustly expressed in the developing brain, with cerebellar expression primarily in PCs (Fig. 1 B, B’, C, C’, green arrowheads). Cre-dependent recombination is efficient and specific at birth, as indicated by TOM expression restricted to cerebellar PCs (Fig. 1 B”, C”, red arrowheads). We confirmed the specificity of Pcp2^Cre^ recombination in the adult mouse brain (Fig. 1 D, D’) in agreement with our prior characterization of this Pcp2^Cre^ mouse line [36].

### Daglα KOs make fewer missteps crossing the horizontal ladder and exhibit normal forelimb coordination

We evaluated motor coordination and forelimb skilled movement in young adult (two-month-old) PC Daglα KO mice. The precision of paw placement was assessed in mice crossing a horizontal ladder with unevenly spaced rungs (Fig. 2 A). Misses (when the paw slips between or off the rung, resulting in the loss of posture) were scored for hind paw placement, revealing a significantly lower percentage of misses in KOs (24 LMs mean = 16.95 %, 23 KOs mean = 10.43 %, the p-value of unpaired T-test = 0.0091, the difference between means ± SEM = −6.520 ± 2.393) (Fig. 2 B). The total time to cross the ladder did not differ between genotypes (23 LMs mean = 22.61 seconds, 21 KOs mean = 22.33 seconds, the p-value of unpaired T-test = 0.9545, the difference between means ± SEM = −0.2754 ± 4.793) (Fig. 2 C). Fewer missteps on the horizontal ladder suggest increased ongoing adjustments of paw placement in PC Daglα KOs – possibly linking reduced 2-AG signaling in the cerebellum to increased cerebellar output.

**Figure 2.**
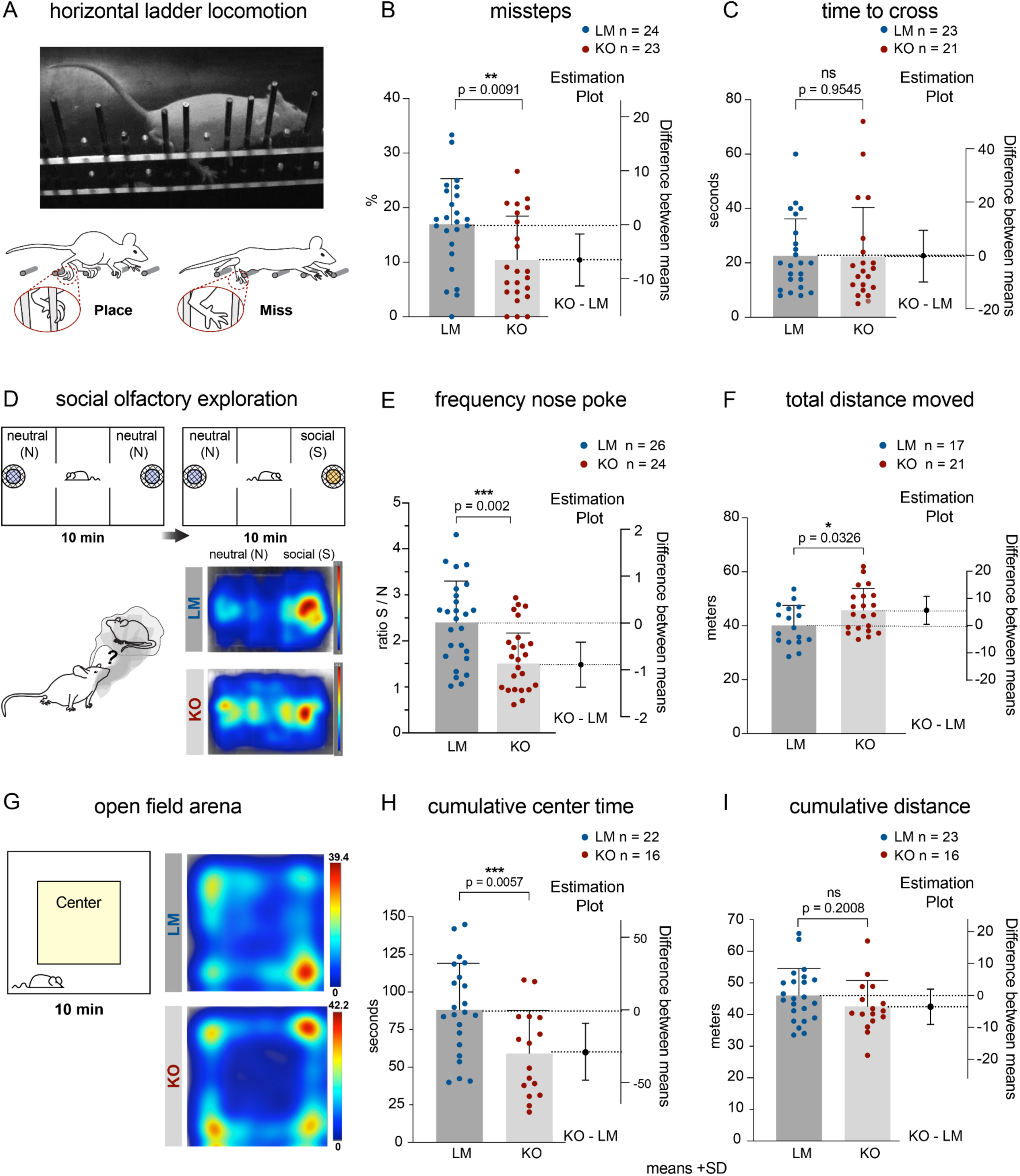
Daglα KOs exhibit increased accuracy of paw placement, decreased social preference, and increased anxiety. (A) The precision of paw placement was assessed in mice crossing a horizontal ladder with unevenly spaced rungs. (B) KOs had a significantly lower percentage of missteps. (C) KOs and LMs took similar times to cross the ladder. (D) Social preference was assessed in an olfactory choice test. Representative heatmaps from a LM and a KO show the LM’s preference to explore the social cues, while the KO exhibits less preference. (E) The preference to explore the social cue (as assessed by the frequency of social over neutral nose pokes) is significantly lower in KOs. (F) KOs exhibit increased activity, i.e., longer total distance moved, during the social olfactory exploration test. (G) Representative heatmaps show that KOs exhibit increased avoidance of the center in the open field arena as compared to LMs. (H) The cumulative time in the center of the open field arena is significantly lower in KOs. (I) The cumulative distance traveled during 10 minutes of exploring the open field does not differ between the genotypes. Columns show means + SD. An unpaired student T-test was used to assess the p-values.

Learning and coordination for the execution of skilled forelimb movements were assessed in the millet seed reaching and retrieval task. In this task, the mice learn a new skill – to reach through a narrow slit to retrieve a millet seed. To evaluate hand dexterity, the percentage of successfully retrieved seeds was assessed for ten consecutive days after the first three days of learning the task (Fig. S1 A). After 30 days of rest, the mice were re-introduced to the testing chamber to assess their motor memory of the task. We observed no differences between genotypes in the seed retrieval task or in the ability to perform the task one month later (Fig. S1 B).

### Daglα KOs exhibit reduced social preference and increased anxiety but no changes in repetitive and impulsive behaviors or in predator-smell-induced fear

We conducted a social olfactory preference test, where the mouse was placed in a three-chamber arena with two wire cups covering either clean bedding (neutral) or bedding from a cage of unfamiliar age- and sex-matched mice (social). Representative heatmaps illustrate that a LM spends more time exploring the social side of the chamber, while a KO exhibits less preference for the social side (Fig. 2 D). The preference to explore social cues was assessed by calculating the ratio between the frequency of nose pokes to sniff the social over the neutral cup. LMs exhibit a greater than a two-fold preference for social over neutral cues, but in KOs, the preference is significantly reduced (26 LMs mean = 2.4, 24 KOs mean = 1.6, a p-value of unpaired T-test = 0.002, the difference between means ± SEM = −0.7570 ± 0.2310) (Fig. 2 E). KOs exhibited slightly elevated overall locomotion (17 LMs mean = 40.31 meters, 21 KOs mean = 45.92 meters, a p-value of unpaired T-test = 0.0326, the difference between means ± SEM = 5.609 ± 2.524), suggesting that the reduction in the percentage of social nose pokes is not due to decreased overall activity (Fig. 2 F).

We conducted an open-field test, which indicated increased anxiety in KOs, who spend less time in the center of the arena than LMs (23 LMs mean = 88.18 seconds, 16 KOs mean = 59.21 seconds, a p-value of unpaired T-test = 0.0057, the difference between means ± SEM = −28.97 ± 9.844) (Fig. 2 G and H). We did not see a statistically significant difference between genotypes in the total distance moved in the open-field arena (23 LMs mean = 46.09 meters, 16 KOs mean = 42.54 meters, a p-value of unpaired T-test = 0.2008, the difference between means ± SEM = −3.545 ± 2.722) (Fig. 2 I). To conclude, KOs exhibit reduced social preference and increased anxiety.

Increased anxiety is often associated with altered fear responses. To evaluate predator smell fear induced immobility in KOs, we compared freezing in mice exposed to the smell of water and Trimethylthiazoline (TMT), an aromatic compound in fox urine. Our analysis revealed no differences in fear response between genotypes, suggesting that both predator-induced fear and the sense of smell are normal in KOs (Fig. S1 D).

We assessed grooming and marble burying to evaluate repetitive behaviors. The percentage of time spent grooming out of 20 minutes of video-recorded spontaneous behavior did not differ between the genotypes (Fig. S1 E). Similarly, LMs and KOs did not differ in the percentage of marbles buried (Fig. S1 F). In addition, we evaluated impulsive behaviors by assessing the ratio of reaches in the seed-reaching task when there was no seed on the tray in front of the slit (i.e., impulsive reaches) and again saw no differences between genotypes (Fig. S1 C). Our analysis suggests that cerebellar PC-specific Daglα KOs do not exhibit deficits in repetitive or impulsive behaviors.

We conclude that KOs exhibit no deficits in predator-smell-induced fear, nor in impulsive and repetitive behaviors.

### Overall physical development, anatomy, synaptic distribution, and basal synaptic strength are normal in PC Daglα KOs, while CB1 is downregulated in granule cell axons

Studies on the neurodevelopmental roles of eCB signaling have focused on the fore- and midbrain, where developmental disruptions of eCB signaling result in axon misrouting, aberrant synaptic development, and motor, cognitive, and emotional deficits [37–45]. The contribution of eCB signaling to cerebellar development is not well characterized, but we expected it to play a prominent role based on findings in other brain regions and on our recent work that showed robust expression of the eCB signaling system in the developing cerebellum; and in global CB1 KOs – reduced size, altered morphology of anterior cerebellar vermis, and deficits in forelimb coordination [46]. Utilizing the PC Daglα KOs, where Daglα deletion is induced around birth and is restricted to PCs, allows us to assess the requirements for PC-derived 2-AG in the structural and functional postnatal development of cerebellar circuits. We conducted our anatomical analysis at two months of age, the endpoint of cerebellar neurocircuit development.

In the LMs, Daglα is highly expressed throughout the PC layer (Fig. 3 A, A’). Like other molecular markers of PCs, Pcp2^Cre^ is expressed in periodic domains along the PC layer, and Cre-recombined PCs null for Daglα appear as striped patches in the KO (Fig. 3 B, B’). Higher magnification confocal microscopy with double immunostaining for Daglα (red) and Cre recombinase (green) confirmed the specificity and selectivity of the Daglα KO from the Cre-positive PCs (Fig. 3 C-C”).

**Figure 3.**
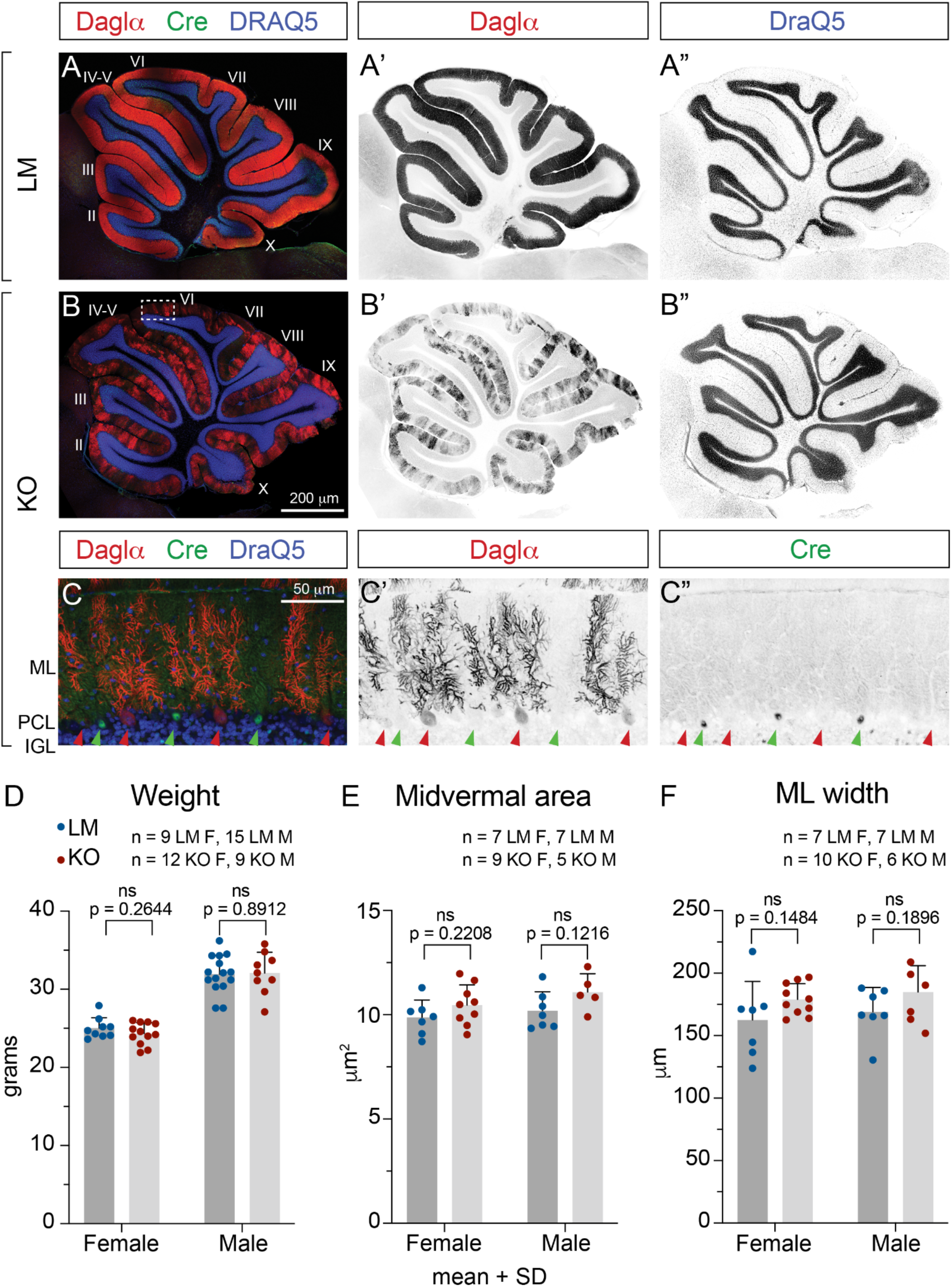
Daglα ablation from cerebellar Purkinje cells does not affect cerebellar gross anatomy. Midsagittal sections from two-month-old cerebella show that (A-A’) Daglα (red) is prominently expressed in the Purkinje cell layer in LMs. (B-B’) Daglα is lost in a subset of PCs in KOs. (A” and B”) Nuclear DRAQ5 labeling (blue in A and B) shows normal cerebellar vermis anatomy. (C-C’’’) A higher magnification view of lobe VI (dotted rectangle in B) shows that Daglα expression is absent specifically in Purkinje cells that express Cre recombinase (green – highlighted by green arrowheads), while Cre-negative Purkinje cells continue to express Daglα, localized primarily to dendrites (ML) and soma (PCL). (D) KO weight is normal at 2-month-old. KOs exhibit no differences in anatomical parameters such as (E) the area of cerebellar midvermis or (F) ML width. ML = molecular layer, PCL = Purkinje cell layer. IGL = inner granule cell layer. Roman numerals designate the lobules of the cerebellar vermis. Columns show means + SD. An unpaired student T-test was used to assess P-values.

Molecular stripe domains are well characterized in PCs for many physiologically important proteins, including Plcβ4 (Fig. S2 B, C, B”, C”). Plcβ4 plays a key role in the signaling cascades downstream from activation of AMPA and mGluR postsynaptic glutamatergic receptors, stimulating Daglα synthesis of 2-AG upon PC depolarization [47]. Immunohistochemistry reveals Daglα expression throughout the PCL, with distinctly different levels of expression in periodic PC clusters (Fig. S2 C, C’). The high Daglα-expressing PCs obey the domain boundaries of the classical molecular stripe domains [48] as identified by Plcβ4 (Fig. S2 A-C”) and localize to Plcβ4-negative stripes (Fig. S2 C-C”). KOs exhibit patches of Daglα-negative PCs in all cerebellar zones (Fig. S2 D, D’, E, E’). However, the pattern of Plcβ4 stripes is unaffected (Fig. S2 E”). These observations suggest that the regulation of Plcβ4 expression in PCs is independent of the levels of eCB signaling controlled by Daglα.

Our analysis shows that overall physical development is normal in PC Daglα KOs: the KOs are indistinguishable from LM controls by body weight at two months old (Fig. 3 D). Anatomical parameters, such as total midvermal area and molecular layer (ML) width, are normal in KOs (Fig. 3 E and F), suggesting that the numbers of cerebellar neurons and their localization into layers are unaffected by loss of Daglα from PCs.

Synaptic competition, retrograde signaling, and neuronal activity regulate the numbers and targeting of excitatory and inhibitory synapses along the developing PC dendrites and somata [36,49–52]. To elucidate whether PC-derived 2-AG plays a role in synaptic placement and the refinement of synaptic territories in the PC somatodendritic compartment, we analyzed the density and distribution of excitatory and inhibitory synapses on PC dendrites and somata at two months old in the anterodorsal domain of midvermal lobe VI (Fig. 4). PC afferents can be distinguished by their specific expression of presynaptic markers: granule cell axons (parallel fibers) express vGluT1 and make synapses on PC dendritic spines; climbing fiber synapses localize to PC dendritic shafts and express vGluT2; molecular layer interneurons express vGaT and make synapses on PC somata and dendritic shafts. We collected 5-μm-thick confocal stacks and used Imaris to generate spots from vGluT1, vGluT2, and vGaT synaptic puncta. To quantify the distribution and density of synaptic puncta on PC dendritic and somatic compartments, we filtered the spots to include only those adjacent to the PC surfaces. PC surfaces were generated based on TOM expression in PCs and were separated into dendritic and somatic regions of interest (ROIs). We found no differences between genotypes in the density or distribution of vGluT1, vGluT2, and vGaT synapses in PC somata and dendrites (Fig. 4), suggesting that the targeting of PC afferent synapses and the overall structural aspects of cerebellar circuitry are normal in Daglα KO PCs.

**Figure 4.**
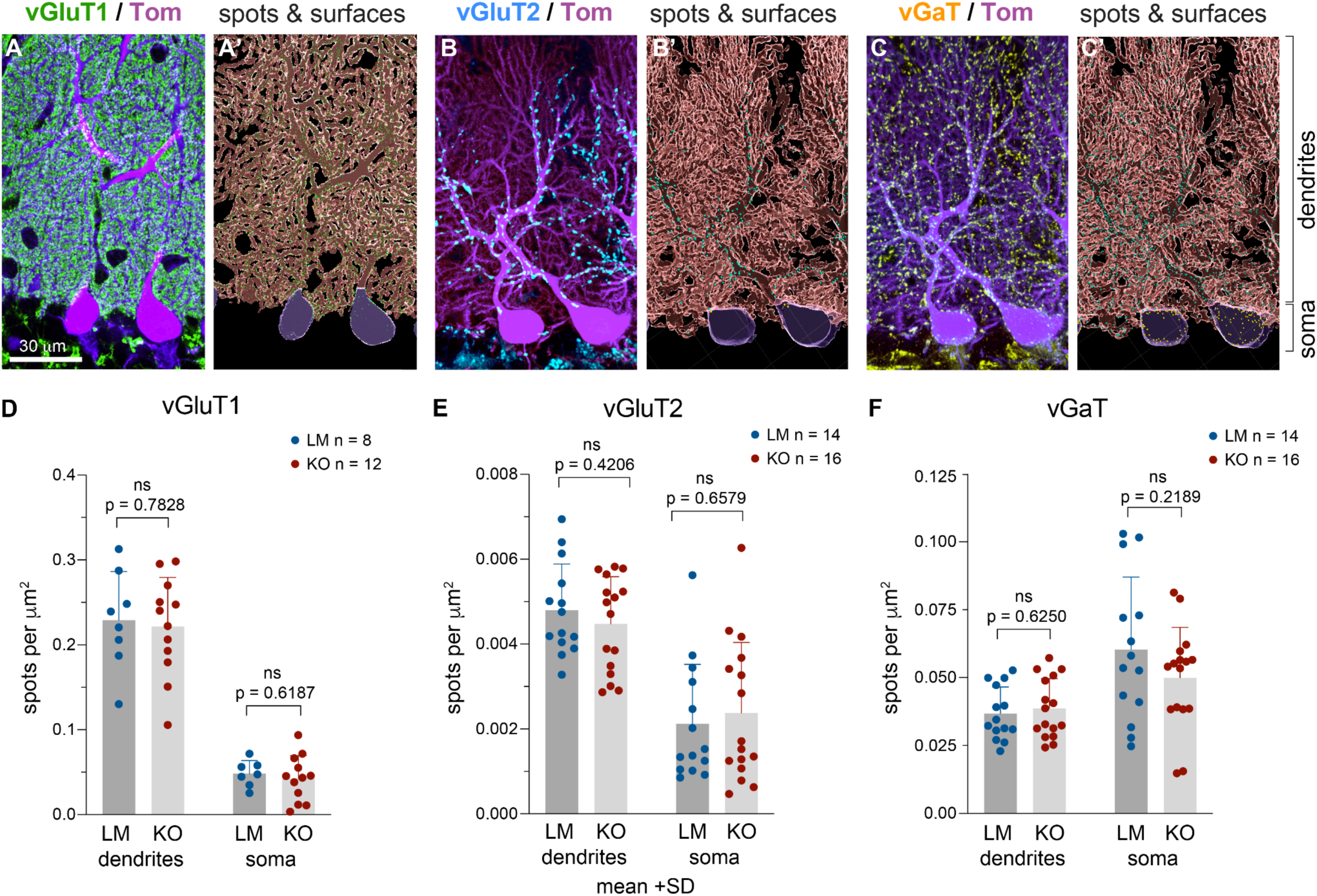
Daglα KO PCs exhibit normal synaptic localization and density. (A-C’) Midsagittal cerebellar vermis anterior-dorsal lobe VI in two-month-old mice. Purkinje cells are identified by TOM expression (purple), GC synapses by vGluT1 (green), CF synapses by vGluT2 (blue), MLI synapses by vGaT (yellow). Imaris was used to reconstruct PC surfaces and the spots of synaptic puncta, filtered to show only those within 0.2 µm of the PC surface, and classified as “on soma” or “on dendrites.” (D-F) Graphs show the density of synaptic spots per area of dendrites and soma. No significant differences in synaptic density or distribution (soma versus dendrite) were found in Daglα KO PCs. Columns show means + SD. An unpaired student T-test was used to assess P-values.

CB1 expression and localization can be regulated by ligand abundance. For example, CB1 expression and synaptic localization are reduced when 2-AG levels are elevated in MAGL KOs [53] or following exposure to exogenous CB1 agonists [54]. Altered CB1 expression can, in turn, affect other types of cannabinoid-dependent synaptic plasticity and neuronal sensitivity to exogenous cannabinoids. Previous studies demonstrated that this signaling-intensity-dependent regulation of CB1 synaptic abundance is particularly robust in excitatory synapses [55]. Our analysis of CB1-positive voxels within vGluT1-positive parallel fiber axons and presynaptic terminals revealed a significant reduction of presynaptic CB1 localization in Daglα KOs (9 LMs mean = 11848 μm^3^, 12 KOs mean = 6474 μm^3^, a p-value of unpaired T-test = 0.0109, the difference between means ± SEM = −5374 ± 1905) (Fig. 5).

**Figure 5.**
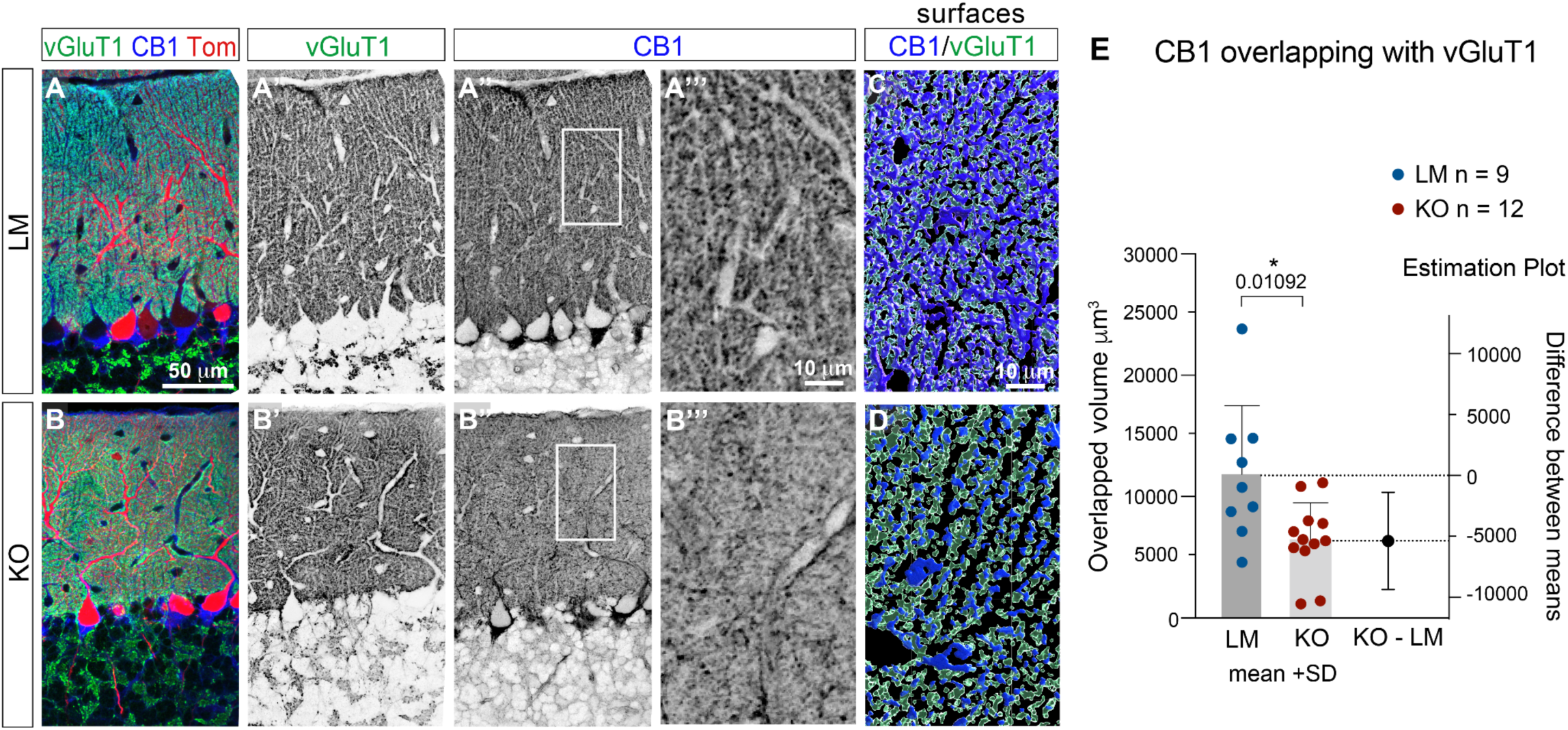
CB1 expression is reduced in vGluT1-positive parallel fiber terminals onto Daglα KO PCs. (A-B’’’) Confocal stacks were collected from the midsagittal cerebellar vermis anterior-dorsal lobe VI in two-month-old mice. (A’’’, B’’’) zoomed-in views of rectangular regions in A” and B”. Purkinje cells are identified by TOM expression (red). vGluT1 (green) and CB1 (blue) expression overlap in parallel fiber synapses in LM (A-A’’’). However, in KOs (B-B’’’) CB1 expression is reduced. (C-D) Imaris was used to construct surfaces of vGluT1 (green) and CB1 (blue) and to assess volume overlap, showing (E) a significant reduction in CB1 overlap with vGluT1 volumes. Columns show means + SD. An unpaired student T-test was used to assess the P-value.

To assess whether synapses terminating onto Daglα KO PCs exhibit any functional alterations in basal synaptic strength, we evaluated spontaneous excitatory and inhibitory postsynaptic potentials (sEPSC and sIPSC) in Daglα KO PCs. The recordings were done from cerebellar slices in control TOM-positive PCs in LMs, and in TOM-positive Daglα KO PCs in KOs (Fig. S3 A). Daglα KO PCs exhibit normal basal synaptic strength in both excitatory and inhibitory synapses as assessed by amplitude, frequency, rise time and decay time (tau) of sEPSCs (Fig. S3 B-B’’’) and sIPSCs (Fig. S3 C-C’’’). In sum, both excitatory and inhibitory synapses on Daglα KO PCs exhibit normal localization and basal functional properties. However, CB1 expression is reduced in vGluT1-positive granule cell axons and presynapses.

### Depolarization-induced short-term synaptic plasticity is dramatically diminished in Daglα KO PCs

Concurrence of synaptic inputs onto PCs controls patterns of PC activity and, consequently, cerebellar-influenced behaviors [56]. Retrograde eCB signaling from PCs regulates the strength of all these synapses: PC depolarization triggers an elevation in cytoplasmic calcium, leading to the release of 2-AG that retrogradely activates presynaptic CB1, causing reduced neurotransmitter release from both excitatory synapses (i.e., depolarization-induced suppression of excitation, DSE [12]) and inhibitory synapses (depolarization-induced suppression of inhibition, DSI [6]). Both DSE and DSI recover to baseline after about one minute. This timescale is consistent with on-the-fly adjustments of ongoing behaviors likely to regulate the accuracy of paw placement or the persistence in the exploration of social cues. Using global Daglα KOs, DSE and DSI were shown to be dependent on Daglα expression [9] but PC-specific requirement for Daglα in DSE and DSI has not been investigated before.

We assessed DSE and DSI in cerebellar slices by measuring evoked excitatory and inhibitory postsynaptic currents (eEPSCs and eIPSCs) in PCs after stimulating parallel fibers and comparing eEPSCs and eIPSCs amplitudes before and after PC depolarization. LMs exhibited more than two-fold reduction in eEPSC and eIPSC amplitudes directly following PC depolarization, exhibiting robust DSE and DSI, but in KOs, post-depolarization, eEPSC and eIPSC amplitudes were very similar to pre-depolarization (Fig. 6 A, D), and returned to baseline faster (Fig. 6 B, E). The ratio of cumulative amplitudes 30 seconds before versus 30 seconds after depolarization shows a reduction to less than half of eEPSC and eIPSC amplitudes in LMs, but only to ~20% of pre-depolarization eEPSC amplitude (Fig. 6 C), and ~10% of eIPSC amplitude (Fig. 6 F) (DSE cumulative amplitude ratio: 7 LMs mean = 0.3796, 9 KOs mean = 0.7836, p-value of unpaired T-test = 0.0031, the difference between means ± SEM = 0.4040 ± 0.1131; DSI cumulative amplitude ratio: 4 LMs mean = 0.5057, 9 KOs mean = 0.9149, p-value of unpaired T-test = 0.0158, the difference between means ± SEM = 0.4092 ± 0.1435). Supplemental Figure 5 shows eEPSCs and eIPSCs before and after PC stimulation in KOs and LMs without normalization (Fig. S5 A, B). In conclusion, both DSE and DSI are nearly abolished in Daglα KO PCs, confirming the requirement for Daglα expression in PCs and retrograde 2-AG signaling for these forms of short-term synaptic plasticity.

**Figure 6.**
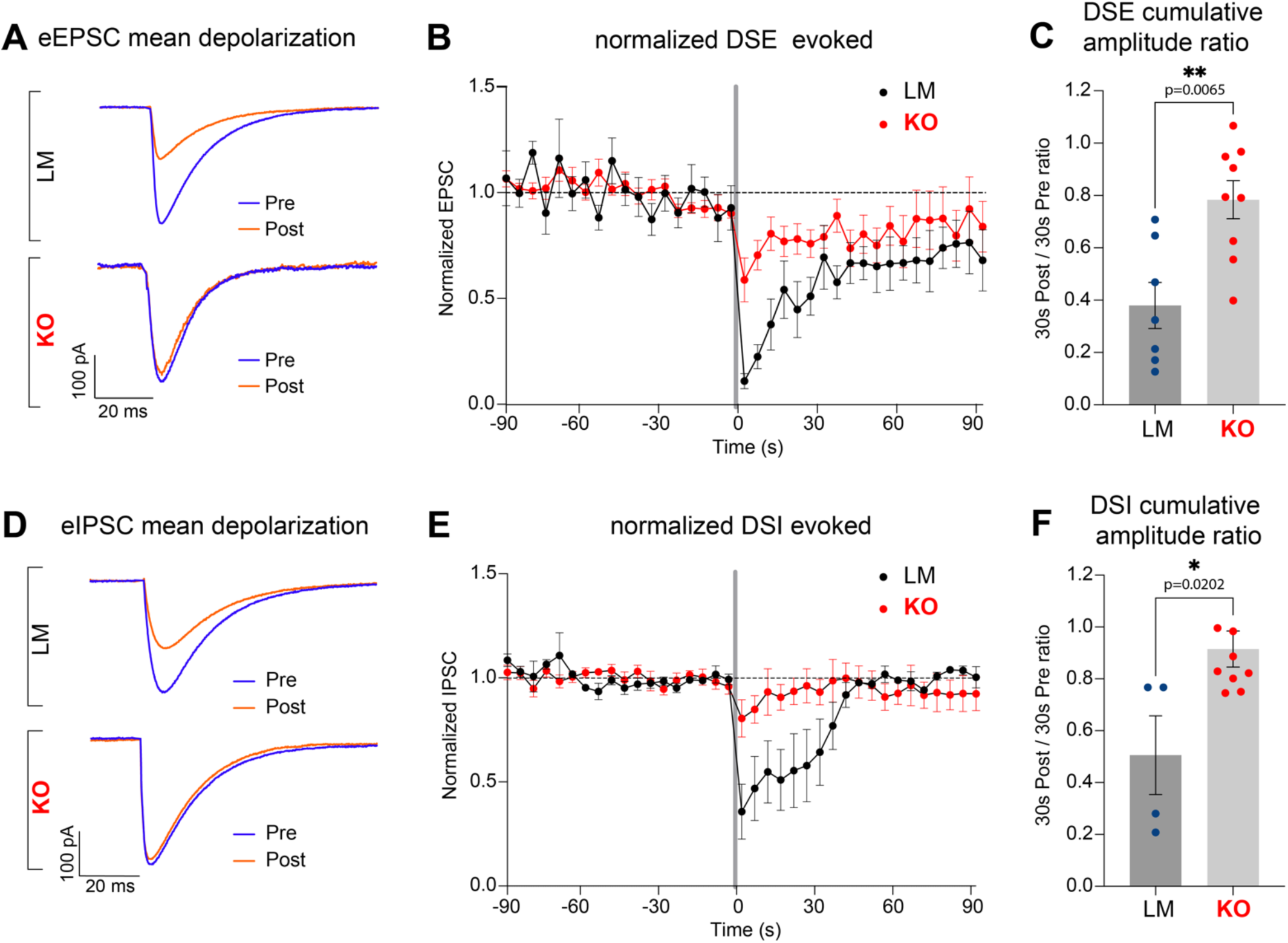
Short-term synaptic plasticity, DSE and DSI, is diminished in Daglα KO Purkinje cells. (A) Representative evoked excitatory postsynaptic currents (eEPSC) before (blue = pre) and after (red = post) Purkinje cell depolarization. LM exhibits smaller post-than pre-eEPSC amplitudes, a hallmark of depolarization-induced suppression of excitation (DSE). However, KOs exhibit almost no difference in pre- and pos-eEPSC amplitudes, i.e., DSE is strongly reduced in the KO. (B) Timecourse of the changes in eEPSC amplitudes before and after Purkinje cell depolarization. In LMs, eEPSC amplitudes are reduced following Purkinje cell depolarization (Time 0), gradually returning to baseline over the next 90 seconds. In the KOs, the reduction in eEPSC amplitudes following Purkinje cell depolarization is markedly smaller. (C) The aggregated cumulative ratio of eEPSCs from 30 seconds before and 30 seconds after Purkinje cell depolarization. LMs exhibit robust DSE, with the post-amplitude reduced to less than half of the pre-amplitude. KOs exhibit small residual DSE, with the pos-eEPSCs reduced to about 0.8 of the pre. 7 LM and 9 KO cells were recorded. (D) A much smaller inhibitory postsynaptic current (eIPSC) amplitude following Purkinje cell depolarization (red=post compared to blue=pre) indicates depolarization-induced suppression of inhibition (DSI) in LMs. The difference between pre- and post-depolarization amplitudes in the KO is very small. (E) In LMs, timecourse shows robust suppression of eIPSC amplitudes following Purkinje cell depolarization in LMs, with a return to baseline within 60 seconds. In contrast, KOs exhibit very little DSI, with only a small difference between pre- and post-eIPSC amplitudes and a faster return to baseline. (F) The ratio of cumulative eIPSC amplitudes 30 seconds before and 30 seconds after Purkinje cell depolarization, showing that in LMs the post-eIPSCs are about half of the pre, but in KOs there is almost no reduction consistent with dramatically reduced DSI. 4 LM and 9 KO cells were recorded. Columns show means (+-SEM). An unpaired student T-test was used to assess p-values.

### Purkinje cell activity is reduced during social exploration, as indicated by reduced cFos expression in PCs

Due to the recurrent excitatory and inhibitory loops in cerebellar circuits, it is difficult to predict how the combination of greatly reduced DSE and DSI affects PCs’ net activity. We utilized immunodetection of the immediate early gene, cFos, as a marker of recent elevated activity in the cerebellar cortex following the olfactory social choice task. cFos is a transcription co-activator characterized by rapid turnover and robust upregulation in response to cellular activity, resulting in a sharp peak of maximum expression at about one hour after an elevated activity [57]. We focused on vermal lobes VI-VII, which are involved in social and emotional behaviors [58,59]. In LMs one hour after the olfactory social choice test, midvermal lobes VI-VII show abundant cFos-positive cells (Fig. 7 A, A”), as well as robust Daglα expression in PCs (Fig. 7 A’). Zoomed-in views of anterodorsal lobe VI (Fig. 7 B, B”, white rectangle in Fig. 7 A) shows that cFos positive cells are present in all three layers of the cerebellar cortex: the inner granule cell layer (IGL), the molecular cell layer (ML) that contains somata of molecular layer interneurons (MLIs), and the Purkinje cell layer (PCL) that contains the monolayer of PC somata (purple arrowheads highlight Daglα expression PCs, and green arrowheads point to cFos expression in PC nuclei). KOs, as expected, exhibit patches of Daglα-negative PCs (Fig. 7 C’, D’). In KOs, cFos expression in IGL and ML is comparable to LMs, but significantly lower in PCs (Fig. 7 C-D”). The density of cFos-positive PCs normalized to the PCL length in lobes VI-VII is lower in KOs as compared to LMs (Fig. 7 E) (14 LMs mean = 18.08 cells per 100 μm, 16 KOs mean = 13.39 cells per 100 μm, a p-value of unpaired T-test = 0.0214, the difference between means ± SEM = −4.690 ± 1.924).

**Figure 7.**
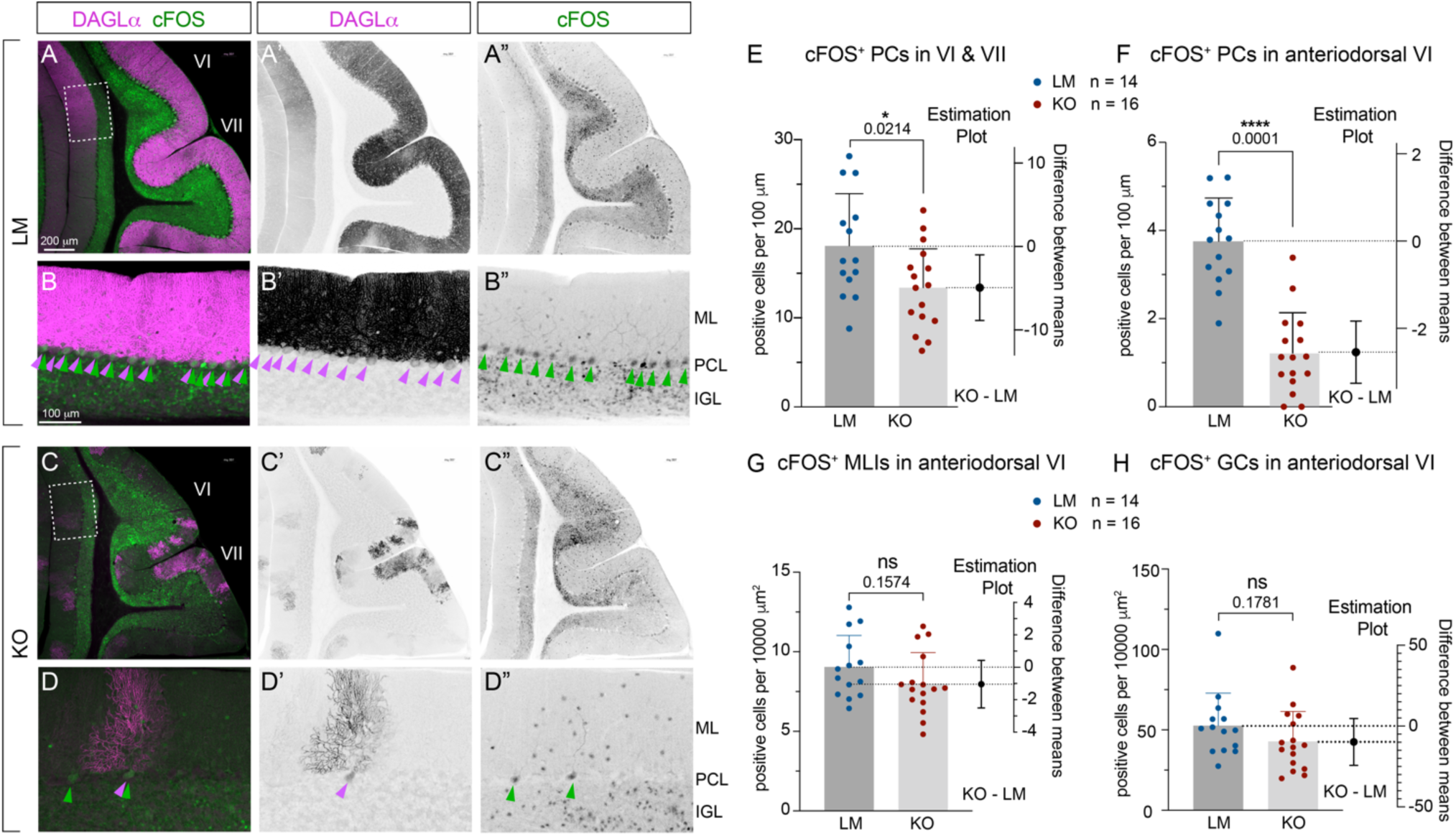
cFos expression is high in Purkinje cells in midvermal lobes VI-VII after the social olfactory exploration task in LMs, but low in KOs. (A-A” and C-C”) lobes VI-VII imaged from sagittal midvermal sections. Daglα expression (purple), which is high in LMs, is absent in a subset of Purkinje cells in KOs. (A-B”) In LMs following social exploration cFos expression (green) is high in all three layers: granule cells (GCs), Purkinje cells (PCs), and molecular layer interneurons (MLIs). (B-B,” D-D,”) Higher magnification images from anterior-dorsal lobe VI (dotted rectangles in A and C). Daglα^+^ PCs are marked by purple arrowheads, and cFos^+^ PC nuclei are highlighted by green arrowheads. (C-D’’) cFos expression is reduced in most PCs but remains high in GCs and MLIs. (E) Quantification shows that cFos-positive PCs per length of the Purkinje cell layer in lobes VI-VII are reduced in the KOs. (F) The anterior-dorsal portion of lobe VI (highlighted by dotted squares in A and C) exhibit the most dramatic reduction in cFos-positive PCs in the Daglα-KOs. (G and H) No statistically significant differences in cFos expression in GCs and MLIs between genotypes. Columns show means + SD. An unpaired student T-test was used to assess P-values.

The midvermal region of lobes VI-VII is likely to contain many distinct functional zones, with distinct patterns of cFos activation. For consistency between animals, we repeated the analysis focusing on a small anatomical area with well-defined landmarks, the anterodorsal region of lobe VI (dotted rectangles in Fig. 7 A and C). In this region, cFos expression in PC is much lower in KOs as compared to LMs (Fig. 7 F), but cFos expression in the IGL (Fig. 7 G) and the ML (Fig. 7 H) is similar between the genotypes. In conclusion, Daglα KO PCs exhibit reduced activity during social olfactory exploration, which correlates with reduced preference to explore social cues and diminished short-term synaptic plasticity.

## DISCUSSION

### PC Daglα KO mouse model highlights the role of cerebellar eCB signaling in the regulation of social behaviors and anxiety

We generated a PC-specific Daglα KO mouse line (PC Daglα KO) to examine how the reduction in PC-derived 2-AG affects the development and function of cerebellar circuits and to evaluate the contribution of the cerebellar eCB signaling to ASD-associated behavioral pathology. The results of this study support the key role of PC-derived 2-AG signaling in the regulation of social approach and anxiety, linking cerebellar PC activity levels and short-term synaptic plasticity to the regulation of social behaviors. Our findings highlight the importance of further investigation into the mechanisms that regulate PC activity as potential therapeutic targets for treatments of neurological disorders associated with reduced sociability and increased anxiety.

The molecular and phenotypic heterogeneity in individuals with ASD is staggering. Adding to the complexity, not only have mutations in hundreds of different genes been identified that predispose humans to ASD (and monogenic ASD-like disorders), but many of these mutations and diagnostic criteria overlap with other neurodevelopmental disorders [60]. To facilitate a deeper understanding of neurodevelopmental mechanisms underlying these disorders, a focus on the molecular deficits associated with narrowly defined behavioral domains and their underlying neural circuitry can be very useful [61,62]. Consequently, transgenic mouse models harboring mutations in specific genes associated with monogenic ASD-like neurodevelopmental disorders have been generated and have been instrumental in improving the understanding of disease pathology. Yet, as ASD manifestations differ between individuals, different mouse models recapitulate different phenotypes, and generating new models is essential to elucidate cellular, molecular, and behavioral endophenotypes not yet addressed by the existing models.

The eCB signaling system is widely expressed in the brain and plays a key role in the regulation of social and emotional behaviors as demonstrated by reduced social preference and increased anxiety in global Daglα KO mice [22,23]. The role of 2-AG in the regulation of anxiety has been specifically investigated in the striatum, where cell-type-specific Daglα KO in dopamine receptor 1 expressing medium spiny interneurons (dMSIs) recapitulates decreased social preference and increased anxiety of the global Daglα KOs [24]. Past studies suggest that eCB neuromodulation contributes to the regulation of emotional states through the modulation of activity in the hippocampal-amygdalar-limbic axis [63].

Recent work demonstrated direct monosynaptic cerebellar regulation of the reward circuits in the ventral tegmental area (VTA) [64] and the contribution of these connections to the regulation of social behaviors [65,66]. In addition, polysynaptic but robust functional connectivity has been described between the cerebellum the hippocampus, the amygdala [67], the medial prefrontal, and the inferior parietal cortex [2–4,68,69], areas strongly implicated in the social and cognitive functions associated with ASD. Our results showing that cerebellar PC specific Daglα KO decreases social preference and increases anxiety highlight the role of the cerebellum as an important modulator of social behaviors and emotional states, and the importance of cerebellar neuromodulation in regulating its influence on the hippocampal-amygdalar-limbic axis.

### Hypoactivity of PCs in the posterior cerebellar vermis underlies the behavioral phenotypes in PC Daglα Kos

Functional brain imaging in humans identified increased activation of posterior cerebellar vermis during social and emotional tasks [3], and lesions in that zone are associated with social and emotional deficits [2,67]. In mice, chemogenetic activation or suppression of cerebellar circuits in the posterior vermis alters social interactions [66,67]. Reinforcing the involvement of this cerebellar zone in the regulation of social behaviors, our study shows reduced cFos expression in PCs in lobes VI-VII during social exploration in PC Daglα KOs, highlighting the correlation between PC hypoactivity and reduced preference to explore social cues.

According to the governing hypothesis, the diverse ASD-associated molecular and cellular abnormalities converge on a common pathological mechanism underpinned by the deregulation of the excitatory-inhibitory balance in neural circuits [70,71]. Functional brain imaging in ASD individuals and animal models has predominantly focused on cortical and subcortical regions, highlighting hyperexcitability in the amygdala, frontal, and parietal cortex as ASD hallmarks [72]. However, differences in the local configurations of microcircuits could lead to the opposite effect of the same molecular changes, suppressing, rather than increasing, excitability in other brain regions. Brain regions influence each other’s activity, and elevated excitability of one brain region can cause increased excitation or increased inhibition in a distant brain region. In contrast to increased activity in the cerebral cortex, ASD diagnosis is associated with decreased activity in the cerebellar cortex (reviewed in [28]). Adding to prior research, the results of our study strongly suggest that the predominant contribution of cerebellar dysfunction to reduced sociability stems from the reduced activity of PCs during social exploration.

Reduced activity of PCs enhances cerebellar output [73], suggesting that the reduced activity of cerebellar PCs can lead to a lower threshold for the cerebellum to generate error messages, causing more frequent adjustments of ongoing behaviors and more switches between behavioral programs - such as increased adjustments of paw placement during horizontal ladder locomotion, and increase switching in the exploration between social and neutral cues. This scenario suggests that reduced PC activity in the cerebellar cortex is likely to correspond to increased excitability in the cerebral cortex. Therefore, increasing PC activity in the cerebellum could be a potential future clinical intervention strategy aimed at decreasing cortical hyperexcitability in the context of the relevant neurological disorders.

### The development of cerebellar circuitry is unaffected in PC Daglα Kos

A body of research implicates eCB signaling in the regulation of axon growth [37,74–76] and synaptic maturation [77,78], yet, the role of cerebellar Daglα in the structural development of cerebellar circuits has not previously been investigated before. The bulk of structural and functional development of cerebellar circuits occurs postnatally. In mice, targeting and refinement of synaptic contacts from granule cells [79], mossy fibers [36], climbing fibers [80], and molecular layer interneurons [81] progresses during the first postnatal month. By ~P30, around the same time when PC dendritic morphology and firing patterns also acquire mature characteristics [82,83], the refinement of synaptic targeting territories of PC afferents and interneurons is also complete; and the molecular stripe domains in PCs assume their mature patterns [82,83]. These aspects of developmental circuit refinement are dependent on PC activity [84–87], and are regulated by retrograde signaling from PCs [36,49,52,88,89]. The eCB signaling system is expressed in the developing cerebellum [46]. Since Daglα-regulated 2-AG production is activity-dependent, it is tempting to hypothesize that PC-derived 2-AG could be an activity-dependent retrograde signal that orchestrates the refinement of cerebellar circuits.

However, in this study, we did not find structural deficits in the postnatal development of cerebellar circuits in PC Daglα KOs: cerebellar midvermal area, localization and density of excitatory and inhibitory synapses onto PCs, and spontaneous synaptic activity are normal in PC Daglα KOs, suggesting that cerebellar circuitry development is intact. Furthermore, PC Daglα KO does not affect PC organization into molecular stripe domains as assessed by Plcβ4 expression, even though the boundaries of stronger versus weaker Daglα expressing PC-clusters do obey Plcβ4 stripe domain boundaries. These results suggest that Daglα expression in PCs is dispensable for the postnatal regulation of the development and structural refinement of cerebellar circuits.

It is important to keep in mind that our experimental design does not allow us to evaluate whether Daglα expression at earlier developmental stages or in cell types other than PCs, may be required for the structural development of cerebellar circuits. The Pcp2 promoter that we used to drive Cre expression turns on just before birth, leaving Daglα expression in PCs intact during embryonic developmental stages, when PCs and their afferents proliferate, differentiate, migrate, and extend axons. In addition, since Pcp2-Cre is not expressed in every PC, it is possible that 2-AG from the neighboring cells could compensate for the reduced 2-AG levels in KOs.

### What cellular mechanisms may contribute to the impaired short-term synaptic plasticity and reduced CB1 expression in vGluT1-positive GC axons in the PC Daglα KOs?

As small lipid molecules, eCBs can diffuse locally within brain regions, influencing the activity of neighboring synapses [90], or systemically, crossing the blood-brain barrier [91]. Acute and chronic stress and pathological conditions, including ASD, are associated with altered systemic levels of eCBs, as measured in plasma or saliva [92,93] – however, it is not clear how systemic levels of eCBs correlate with or affect local eCB concentrations in specific neuronal circuits [91]. If local or systemic diffusion of 2-AG contributes significantly to PC short-term synaptic plasticity, it is likely to mask a reduction in 2-AG signaling due to Daglα KO in a subset (Pcp2-positive) of PCs. Nevertheless, Daglα KO PCs exhibit greatly diminished DSE and DSI, showing that the local activity of Daglα is essential for these forms of short-term synaptic plasticity. The small residual DSE and DSI that can be observed in Daglα KO PCs could be potentially explained by the spill-over of 2-AG from neighboring WT PCs or by the contribution of other eCB synthesizing enzymes.

DSE and DSI are examples of activity-dependent short-term synaptic plasticity, i.e., phasic neuromodulation. In cortical and hippocampal inhibitory neurons, a distinct type of eCB-dependent synaptic inhibition is observed. It is termed tonic inhibition and is characterized by the unmasking of enhanced synaptic strength when eCB signaling is blocked [94]. The baseline eEPSCs and eIPSCs in Daglα KO PCs (before PC stimulation, Fig. S5) appear potentiated compared to LMs, suggesting that Daglα in PCs could also be involved in tonic inhibition of excitatory and inhibitory synapses onto PCs. PC Daglα KOs could be used in future work to evaluate this hypothesis.

The excitatory-inhibitory balance in neural circuits could also be affected through the regulation of intrinsic neuronal excitability, and eCB signaling can regulate overall neuronal activity in an autocrine manner [95]. Intracellular signaling cascades, energy metabolism, and intrinsic functional properties of Daglα KO PCs could be evaluated in the future to assess the contribution of cell-autonomous consequences of Daglα KO and the role these changes play in the regulation of neural circuit excitability and behavioral regulation.

CB1 expression levels and localization are reduced following exposure to exogenous agonists or altered levels of eCB signaling [53][54]. We assessed the localization of CB1 to vGluT1 puncta in excitatory GC axons and found significantly reduced levels of CB1. Reduced synaptic localization of CB1 is likely to enhance overall excitatory neurotransmission from GCs to PCs – potentially contributing to increased tonic strength and diminished short and long-term depression in these synapses. We did not quantitatively evaluate CB1 localization to inhibitory synapses in this study, but we expect that it may be less affected since CB1 expression remains prominent along PC dendritic shafts and horizontal trajectories, which is typical for the axons of MLI inhibitory interneurons, while it is reduced adjacent to PC dendritic spines, where GC axons make their synapses (Fig.5 B’’’)

### How do the results of this study expand our understanding of pathology associated with mutations in DAGLα?

The association between rare single gene mutations in *DAGL*α and ASD led to its ranking in the “high-confidence strong candidate autism risk genes category 2” in the gene scoring module in SFARI (https://gene.sfari.org/database/human-gene/DAGLA). Several gene association studies have linked mutations in *DAGL*α (in most cases arising *de novo* in the patients) to neurodevelopmental disorders [18,19]. In 35 pediatric patients (out of 6,032 probands with neurodevelopmental disorder diagnosis), a significant association was found between rare heterozygote variants in *DAGL*α and ASD and seizure disorders, but not ataxia [19]. This study also found that anxiety is frequently diagnosed in patients with mutations in *DAGL*α, but this association was also high in controls (who have not been diagnosed with a neurodevelopmental disorder) [19]. A recent study focused on nine idiopathic pediatric patients with mutations in *DAGL*α and a specific set of symptoms: ataxia, nystagmus, and developmental delay [18]. The two studies together highlight cognitive, social, and motor developmental phenotypes associated with mutations in *DAGL*α. Our study suggests that the reduction of cerebellar Daglα strongly contributes to the symptoms of reduced social preference and anxiety but does not cause seizures or ataxia in our KO mice at the ages examined.

## CONCLUSIONS

Our results suggest that cerebellar eCB signaling plays a key role in the regulation of cerebellar synaptic plasticity and PC activity, linking deficits in cerebellar eCB signaling to decreased PC activity during social exploration and decreased preference for social cues. These results provide novel evidence for the role of cerebellar eCB signaling in the regulation of social behaviors and the therapeutic potential of eCB signaling augmentation or direct stimulation of PC activity for treatments of disorders associated with decreased interest in social interactions. Conversely, despite the robust expression of Daglα in the developing PCs, postnatal ablation of Daglα from PCs does not affect the development of cerebellar circuits, as assessed by the analysis of cerebellar size, layers, and synaptic localization and density.

## MATERIALS AND METHODS

### Mouse Colony

All mice used in this study were maintained on an outbred CD1 genetic background. The CD1 IGS stock was acquired from Charles River, USA (https://www.criver.com/products-services/find-model/cd-1r-igs-mouse?region=3611) and replenished by purchasing additional CD1 breeders from Charles River once a year. A breeding colony was maintained in the vivarium at Indiana University, Bloomington with a 12-hour light/dark cycle under conditions stipulated by the Institutional Animal Care and Use Committee. The *Daglα* floxed (*Daglα^fl^*) mouse line was generated by the group of Dr. Sachin Patel [34]. The Purkinje cell specific Cre line (*Pcp2^Cre^*) was generated by the group of Noboru Suzuki [35]. Ai9 (B6.Cg-Gt(ROSA)26Sortm9(CAG-tdTomato)Hze/J) tomato (TOM) reporter mouse line was purchased from the Jackson Laboratory (https://www.jax.org/strain/007909). We generated triple transgenic mice by crossing the founder lines, as described in the results (Fig. 1 A). First-degree crosses (siblings, parents, etc.) were always avoided.

### Tissue Preparation

Brain tissue was fixed by transcardial perfusion with 1xPBS followed by 4% PFA. Dissected brains were postfixed in 4% PFA at 4°C overnight, switched to 0.2% sodium azide 1xPBS (PBS-Az), and stored at 4°C until sectioning. The brains were sectioned at 70μm using a Leica VT 1000S vibrating microtome.

### Antibodies and immunohistochemistry

Immunohistochemistry was conducted on free-floating 70μm tissue sections. The tissue sections were washed three times in 1xPBS and incubated in BSA blocking buffer (5% BSA, 1% Triton X-100, 1xPBS). Primary antibodies were applied overnight at 4°C in BSA blocking buffer. The tissue was washed three times in 1x PBS, and secondary antibodies (Alexa Flour 488, 596, or 647 from Jackson ImmunoResearch) were applied at 4°C overnight (1:600 in BSA blocking buffer). DraQ5 (Cell Signaling) was used (at 1:5000 in 1xPBS) to visualize cell nuclei. Streptavidin Alexa 488 (Invitrogen) was used at 1:1000 in BSA blocking buffer to visualize biocytin-filled neurons. Slides were coverslipped using Flouoromount G (SouthernBiotech). We used the following

<u>primary antibodies:</u>

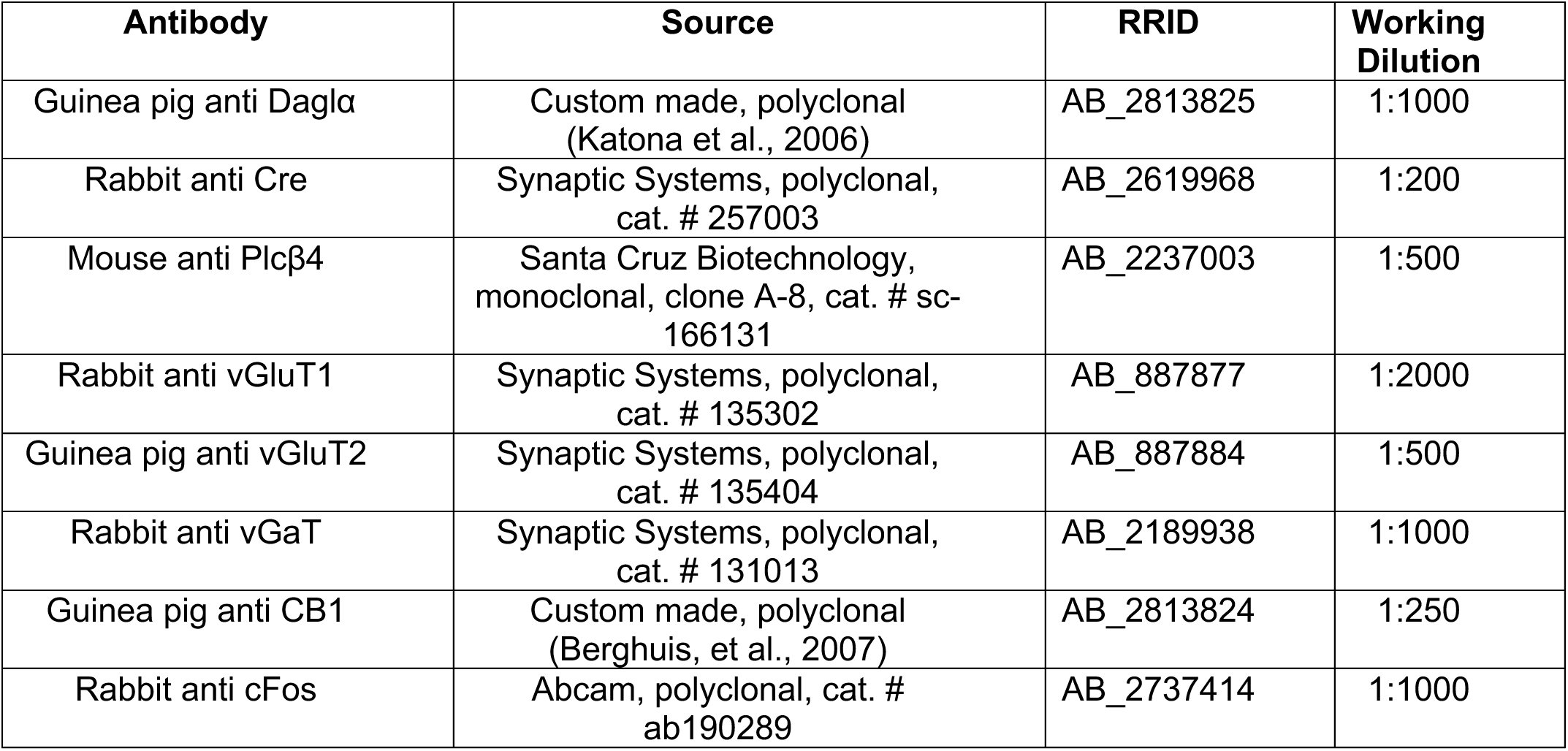

### Microscopy and Image Analysis

A Nikon A1 confocal microscope was used to collect image Z-stacks. The cerebellar area and the layer thickness were analyzed from DraQ5-stained cerebellar sagittal midvermal sections using the free-hand tool to measure areas of interest and distances in FiJi (<u>Fiji</u>). FiJi cell counter tool was used to assess the numbers of cFos-positive cells from max-intensity-projection 5μm-thick confocal image stacks. For the analysis of synaptic density, a 60x objective was used to collect 5μm-thick confocal image stacks from midvermal lobe VI. The density of synaptic puncta per surfaces of Purkinje cell somata and dendrites was assessed in Imaris (Oxford Instruments). Regions of interest were manually outlined for PC somata (PCL) and dendrites (ML). TOM expression in Purkinje cells was used to reconstruct the surfaces of somata and dendrites. Spots were generated based on vGluT1, vGluT2, and vGaT synaptic puncta staining, then filtered to show only those within 0.2 µm of the soma (yellow spots) and dendrite (turquoise spots) surfaces. These spots were classified as “on soma” or “on dendrites.” *<u>Statistical analysis</u>* was performed in GraphPad Prism 10 (www.graphpad.com). For weight, anatomy (midvermal area, ML width) (Fig. 3 D, E, F), synaptic puncta density (Fig. 4 D, E, F), CB1 overlap with vGluT1 (Fig. 5 E), and the density of cFos-positive cells per PCL length/ML&IGL areas (Fig. 7 E, F, G, H) the statistical hypothesis tested the magnitude of differences between KOs and LMs. An unpaired student T-test was used to evaluate means, SEMs, and P-values. Estimation plots were used to analyze the confidence intervals for the differences between means.

### Mouse Behavior

#### Grooming

(Fig. S1 E) Two-month-old mice were individually placed in a plexiglass testing chamber and video recorded for 20 minutes. The videos were manually analyzed for time spent grooming. *<u>Statistical analysis</u>* Statistical analysis was performed in GraphPad Prism 10 (www.graphpad.com). Data was collected from 19 KO and 24 LM animals. The statistical hypothesis tested the magnitude of differences in the percentage of time spent grooming (manually annotated from videos recorded at 60 frames per second) between KOs and LMs. The results are shown as means +SD. P-values were evaluated by unpaired student T-test.

#### Locomotion on the horizontal ladder with uneven rungs

(Fig. 2 B, C) Two-month-old mice were placed on a horizontal ladder with unevenly spaced rungs. The ladder was placed between a clean mouse cage and the test mouse’s home cage, and the mice were videorecorded at 60 frames per second from the ventral side as they walked across the ladder, allowing easy visibility of paw placement. The videos were manually scored frame by frame for the number of missteps and successful placements of hind paws on the rungs. The percentage of missed steps (out of all steps) was calculated. *<u>Statistical analysis</u>* was performed in GraphPad Prism 10 (www.graphpad.com). Data was collected from 23 KO and 24 LM animals. The data is shown as means +SD. The statistical hypothesis tested the magnitude of differences in the percentage of missteps and time to cross the ladder for KOs and LMs. P-values were evaluated by unpaired student T-tests. Estimation plots were used to analyze the confidence intervals for the differences between means.

#### Marble burying

(Fig. S1 F) Two-month-old mice were placed in a clean standard mouse cage filled with 10 cups of corn cob bedding (about 5cm deep) for 15 minutes to habituate. After habituation, the corn cob bedding in the cage was smoothed down, and 8 evenly spaced marbles were placed on top of the corn cob bedding. The mouse was placed back in the test cage and allowed to explore for 15 minutes. After 15 minutes, the number of marbles buried (≤1/4 visible) was scored manually. *<u>Statistical analysis</u>* was performed in GraphPad Prism 10 (www.graphpad.com). Data was collected from 10 KO and 18 LM animals. The data is shown as means +SD. The statistical hypothesis tested the magnitude of differences in the percentage of marbles buried between KOs and LMs. An unpaired student T-test was used to evaluate P-values.

#### Millet seed reaching and retrieval; impulsive reaching

(Fig. S1 A, B, C) To motivate the learning of a new skilled reaching task – extending a forelimb through a narrow slit to reach for, grasp, and retrieve a millet seed – two-month-old mice were food-restricted starting three days before and for the duration of the experiment. The weight of the mice was monitored daily, making sure that no individual lost more than 10% of body weight. Every morning regular mouse chow was added to the cages, estimating the amount of food per day by adding the weight of the chow that corresponds to 10% of the collective weight of the mice in the cage (i.e., if the mice in the cage weighed 100g, 10g of chow was added to the cage once a day). Mice were placed one by one in a plexiglass behavior box from which they reached out through a slit with their forepaw in order to retrieve a millet seed. The experiment started with 3 days of training, during which the mice learned the task. Starting on the fourth day, or after the mice exhibited proficiency in performing the task (being able to retrieve at least 10 seeds in 20 minutes and exhibiting consistent hand preference), the mice were placed in the apparatus for 20 minutes each day for 10 additional days, while seed reaching was video recorded at 60 frames per second. The resulting videos were manually scored for the percentage of successfully retrieved millet seeds out of all the reaching attempts with the seeds present. Millet seeds were added to the tray in front of the slit one by one, and the mice would occasionally reach through the slit even when there were no seeds present on the tray (impulsive reaching). *<u>Statistical analysis</u>* was performed in GraphPad Prism 10 (www.graphpad.com). Data was collected from 7 KO and 9 LM animals. The graph shows % successfully retrieved seeds on each day for a 10-day timecourse. The data is shown as means +SD. Areas under the curve and confidence intervals were assessed for the learning timecourse graphs and showed no differences between KOs and LMs. The percentage of successfully retrieved seeds was evaluated on the last day of testing and one month after the last day of testing. The statistical hypothesis tested the magnitude of differences in the percent of successfully retrieved seeds (out of all attempts where seeds were present on the tray) and the percent of impulsive reaches (out of all reaches) for the last day of testing and the 30-day-later re-testing between KOs and LMs. An unpaired student T-test was used to evaluate P-values.

#### Open field

(Fig. 2 H, I) Two-month-old mice were placed in an open plexiglass arena for 10 minutes. The amount of time spent in the center of the arena and the total distance moved in the arena were calculated. The software used to record the mice during the open field test were Pylon Viewer and EthoVision XT 17. *<u>Statistical analysis</u>* was performed in GraphPad Prism 10 (www.graphpad.com). Data was collected from 16 KO and 23 LM animals. The statistical hypothesis tested the magnitude of differences in cumulative time spent in the center of the arena and cumulative distance explored between KOs and LMs. Estimation plots were used to analyze the confidence intervals for the differences between means and p-values were evaluated by unpaired student T-tests.

#### Predator fear-induced immobility

(Fig. S1 D) Two-month-old mice were individually placed in a plexiglass testing chamber. On day one, the mice were exposed to a cotton ball with 10uL of distilled water (the cotton ball was placed inside a vial suspended out of reach inside the testing chamber). The mice were video recorded at 60 frames per second for 20 minutes. On the next day, the mice were placed in the same test chambers, and video recorded for 20 minutes, but the cotton balls were soaked with 2,5-dihydro-2,4,5-trimethylthiazoline (TMT – an aromatic compound from fox urine) instead of water. The videos were analyzed on the software “EzTrack” for the percentage of time spent immobile. *<u>Statistical analysis</u>* was performed in GraphPad Prism 10 (www.graphpad.com). Data was collected from 20 KO and 24 LM animals. The statistical hypothesis tested the magnitude of differences in time spent freezing between KOs and LMs during exposure to water and TMT. Paired analysis shows significantly increased freezing for TMT compared to water for both genotypes, but there were no differences between the genotypes. P-values were evaluated by 2-way AVOVA.

#### Social olfactory exploration

(Fig. 2 E, F) Two-month-old mice were individually placed in a three-chamber plexiglass arena with two wire cups containing neutral (clean) bedding dishes for 10 minutes to habituate to the arena. After 10 minutes of habituation, one dish was replaced with a social bedding dish (soiled bedding from a cage of sex- and age-matched unfamiliar mice, and the mice explored the arena for another 10 minutes while being video recorded. The software used was Pylon Viewer and EthoVision XT 17. The frequency of nose pokes (to sniff) at the neutral and the social cups was scored manually, and the ratio of social versus neutral nose pokes was calculated. The total distance moved was assessed automatically in EthoVision XT 17. *<u>Statistical analysis</u>* was performed in GraphPad Prism 10 (www.graphpad.com). Data was collected from 24 KOs and 26 LMs for social/neutral nose pokes, and 21 KOs and 17 LMs for the total distance moved since shadows in the arena interfered with automatic tracking for a few animals. The statistical hypothesis tested the magnitude of differences in the frequency of social nose pokes over neutral nose pokes and the total distance moved between KOs and LMs. Estimation plots were used to analyze the confidence intervals for the difference between means. P-values were evaluated by unpaired T-tests.

#### Electrophysiology

(Fig. S4, S5, 6) Postnatal day 25-31 mice were anesthetized with isoflurane. The brains were carefully and promptly removed and blocked. Parasagittal 280μm-thick slices of the cerebellum were cut at 0.1 mm/s with a Leica VT1200 vibratome in ice-cold oxygenated external solution containing a sucrose-based artificial cerebrospinal fluid (aCSF) (in mM: Sucrose, 194; NaCl, 30; NaHCO_3_, 26; Glucose, 10; KCl, 4.5; NaH_2_PO_4_, 0.5; MgCl_2,_ 1; pH 7.4) bubbled with 95% O_2_/5% CO_2_. After cutting, slices were transferred to an incubation chamber containing oxygenated aCSF solution at 33°C for one hour. Slices were then kept at room temperature until recording.

Whole-cell patch recordings were done from Purkinje cells identified by TOM fluorescence at 30-32°C in a submersion chamber perfused (~2ml/min) with aCSF solution comprised of (in mM): NaCl, 124; KCl 4.5; NaH_2_PO_4_, 1.2; MgCl_2_, 1; CaCl_2_, 2; NaHCO_3_, 26; and Glucose 10, continuously bubbled with 95% O_2_/5% CO_2_; pH 7.4, 310 mOsm. Patch pipettes (2–3 MΩ) were pulled from borosilicate glass and filled with internal solution containing (in mM): 120 CsMeSO_3_; 5 NaCl, 10 TEA, 10 HEPES, 5 lidocaine bromide, 1.1 EGTA, 4 Mg-ATP, 0.3 Na-GTP and 0.2% biocytin (pH 7.3, 292 mOsm) for voltage clamp recordings. Biocytin (2 mg/ml) was also added to the internal solution to allow for post hoc visualization of recorded neurons. TOM-positive neurons were identified and visualized with a 40X water-immersion objective on an upright fluorescent microscope (BX51WI; Olympus) equipped with infrared-differential interference contrast video microscopy and epifluorescence through the light path of the microscope using an ultrahigh-powered light-emitting diode (CoolLED). Data were collected using Clampex software (version 10, Molecular Devices). Signals were filtered using a Bessel filter set at 4 KHz and digitized at 50 KHz with a Digidata 1440A A/D interface. A 5mV pulse was delivered regularly to monitor series resistance. Recordings with series resistance greater than 25 MΩ or with changes in access greater than 20% were discarded. For quantification, the software pClamp11(Molecular Devices) and MiniAnalysis (Synaptosoft) were used. Statistical analysis was done using Prism 10 (GraphPad Software). The holding membrane potential was set to −70 mV in all experiments.

Spontaneous excitatory postsynaptic currents (sEPSC) were recorded after adding picrotoxin (50 µM) to the aCSF to block inhibitory currents (9 CONT PCs, 8 KO PCs). Spontaneous inhibitory postsynaptic currents (sIPSC) were recorded after adding NBQX (10 µM) and D-AP5 (50 µM) to block excitatory currents (9 CONT PCs, 8 KO PCs). Amplitude, Frequency, rise time and decay time (tau) were measured. The holding membrane potential was (V_h_) at −70 mV (9 CONT PCs, 8 KO PCs). For evoked PSCs, a stimulating bipolar electrode was placed in the molecular layer, and synaptic responses were evoked at 0.2 Hz (rectangular pulses 0.2 ms duration, 50-300μA).

To induce DSE and DSI, a depolarizing pulse from −70 to 0 mV was applied once for 5 seconds after 2 minutes of baseline recording. Excitatory and inhibitory currents were isolated, as was done in sEPSC and sIPSC recordings described above.

## Author Contributions

Gabriella Smith performed behavior and immunohistochemistry experiments, processed and analyzed data, co-wrote the first draft, and edited the manuscript

Kathleen McCoy performed behavior and immunohistochemistry experiments, designed image analysis protocols, processed and analyzed data, edited the manuscript

Gonzalo Viana Di Prisco performed slice electrophysiology experiments, designed experimental protocols, processed, and analyzed data

Alexander Kuklish performed behavior and immunohistochemistry experiments, analyzed data

Emma Grant performed immunohistochemistry experiments, analyzed data

Mayil Bhat performed mouse behavior experiments

Sachin Patel generated floxed DAGLα mouse line

Brady Atwood designed and directed experiments for slice electrophysiology and interpreted results

Ken Mackie advised in the design of experiments and interpretation of experimental results, edited the manuscript

Anna Kalinovsky designed experiments, directed the project, secured funding, conducted experiments, analyzed data, wrote and edited the manuscript

All authors discussed the results and commented on the manuscript.

## Acknowledgments

We are grateful to Laszlo Barna and Istvan Katona for administering and consulting on the use of the Neuroscience Core Imaging Facility. The imaging was supported in part by P30 grant from NIH - NIDA (P30DA056410). We thank talented high school students Max Rose and Layla Vamos for their assistance in quantifying mouse behavior. We also thank all our lab members for discussing this manuscript. This work was supported by R21 from NIDA DA044000, OVPR-FRSP-SEED and OVPR-FRSP-BRIDG from Indiana University, and CTSI-CORE-PILOT to Anna Kalinovsky, and MH107435 to Sachin Patel.

---

**Supplemental Figure 1.**
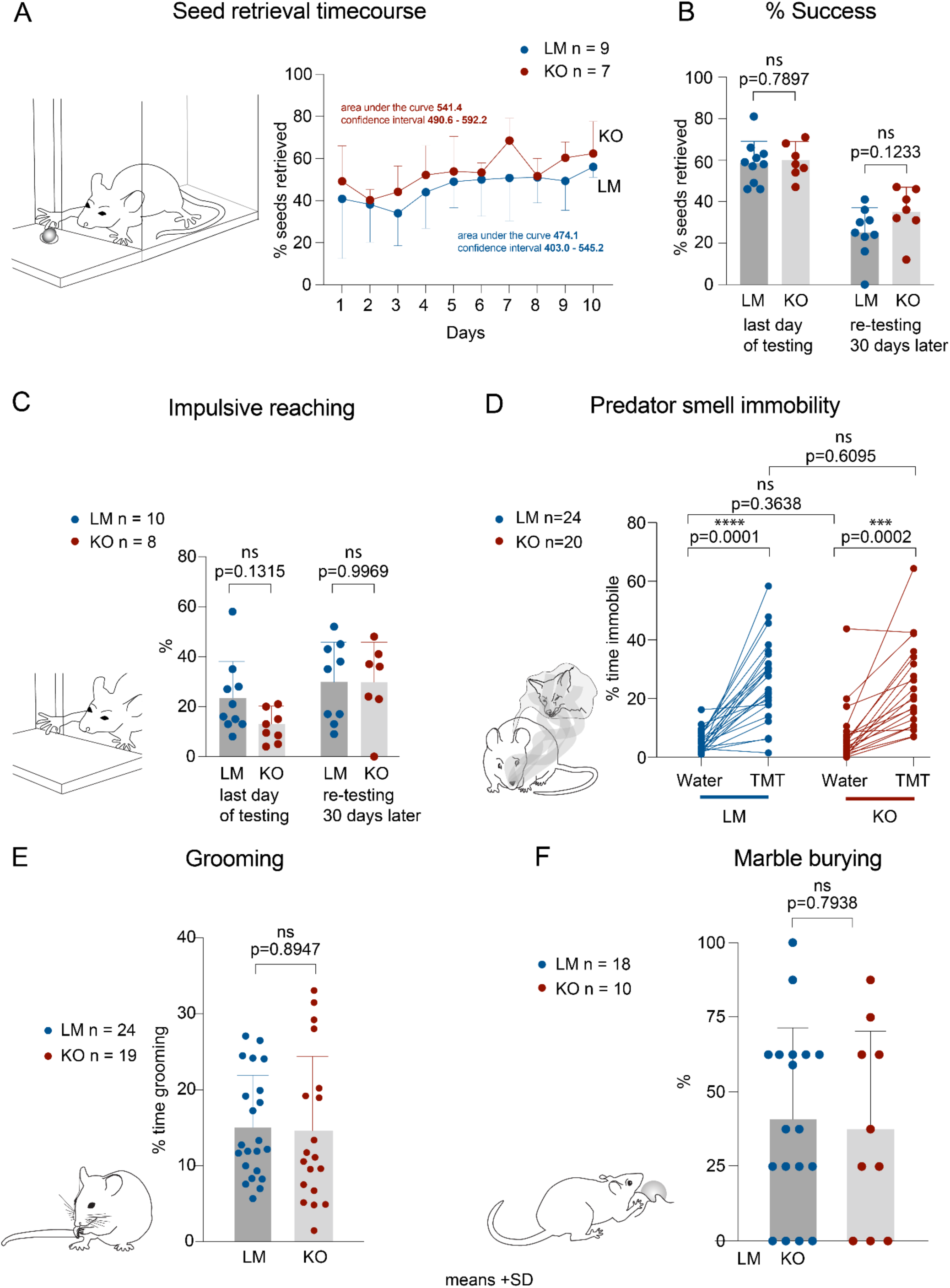
Forelimb skilled movement, predator fear response, impulsive and repetitive behaviors are unaffected in Daglα KOs. (A) The mice were trained to use their forepaws to reach for and retrieve millet seeds through a narrow slit. Their performance was assessed for the percentage of successful reaches over a period of 10 days, revealing no differences between the genotypes in the learning curves. (B) There was no difference between the genotypes on the last day of training or in motor memory retention one month later. (C) Impulsivity was assessed in the seed reaching assay by scoring the percentage of impulsive (reaching when there are no seeds on the tray) out of all reaches. The analysis shows no differences between the genotypes. (D) KOs exhibit normal fear response as assessed by immobility precipitated by exposure to predator smell. (E) Grooming and (F) marble burying were used to assess repetitive behaviors, revealing no differences between the genotypes. Columns show means +SD. An unpaired student T-test was used to assess P-values.

**Supplemental Figure 2.**
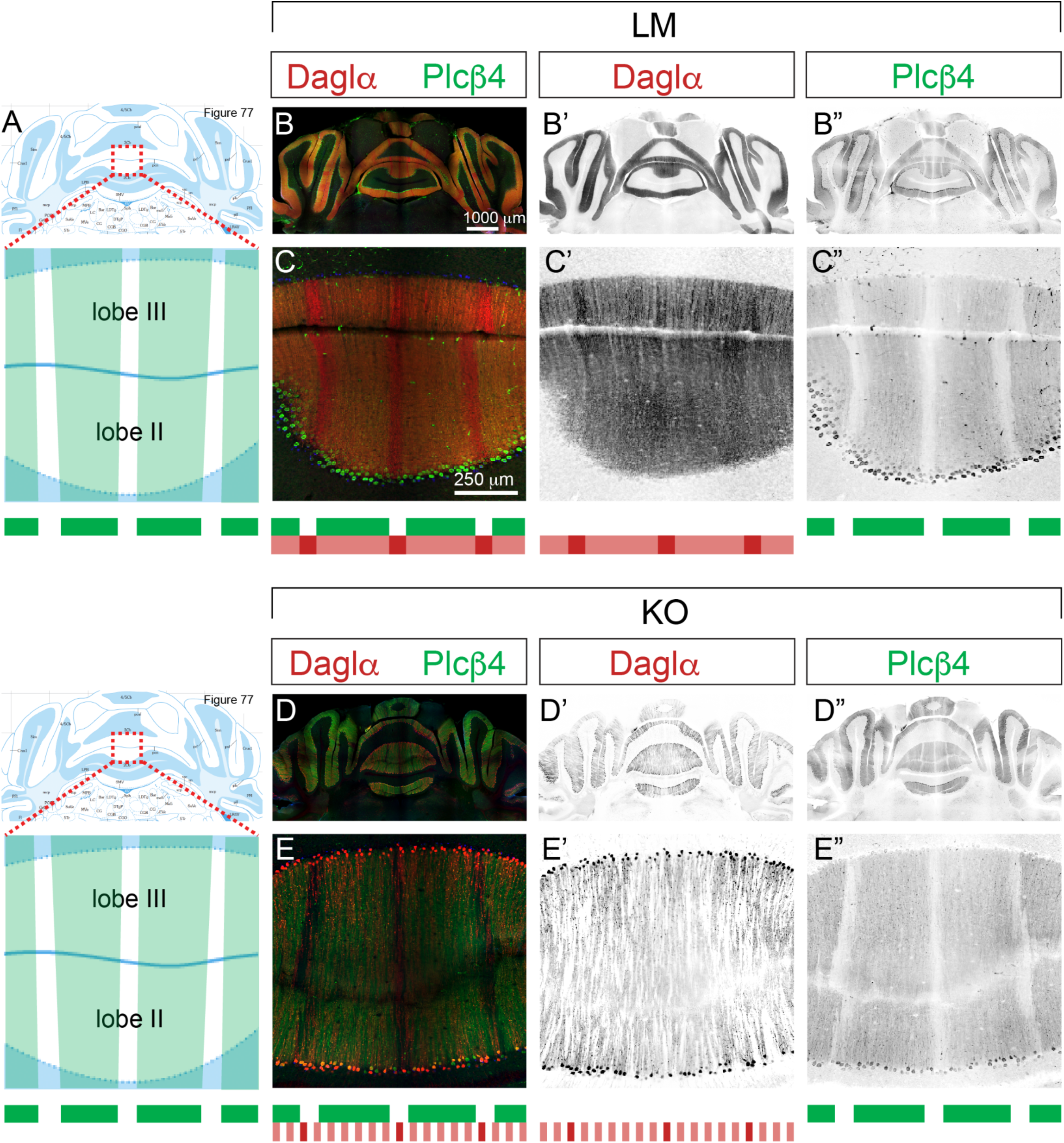
Molecular stripe domains are normal in Daglα KOs. Coronal sections through the anterior zone of cerebellar vermis (lobes II and III) at 2-month-old. (A) Figure 77 from Franklin & Paxinos, “The Mouse Brain in Stereotaxic Coordinates,” shows the plane corresponding to the sections analyzed from LMs and KOs, and in the zoomed-in panel Plcβ4-positive stripe domains are marked with green mask. (B-B’) Daglα (red) is expressed throughout the Purkinje cell layer in all cerebellar zones and lobes and (B”) Plcβ4 (green) expression exhibits the typical alternation of positive and negative stripes in the anterior zone. (C-C’) Higher magnification of lobes II-III showing that Daglα expression exhibits stripe-dependent differences in expression levels: (C”) Plcβ4-negative PCs express higher levels of Daglα (summarized in the stripe diagrams below). (D, D’, E, E’) Patches of Daglα-negative PCs are evident in all cerebellar zones in KOs. However, (E”) the pattern of Plcβ4 stripes is unaffected.

**Supplemental Figure 3.**
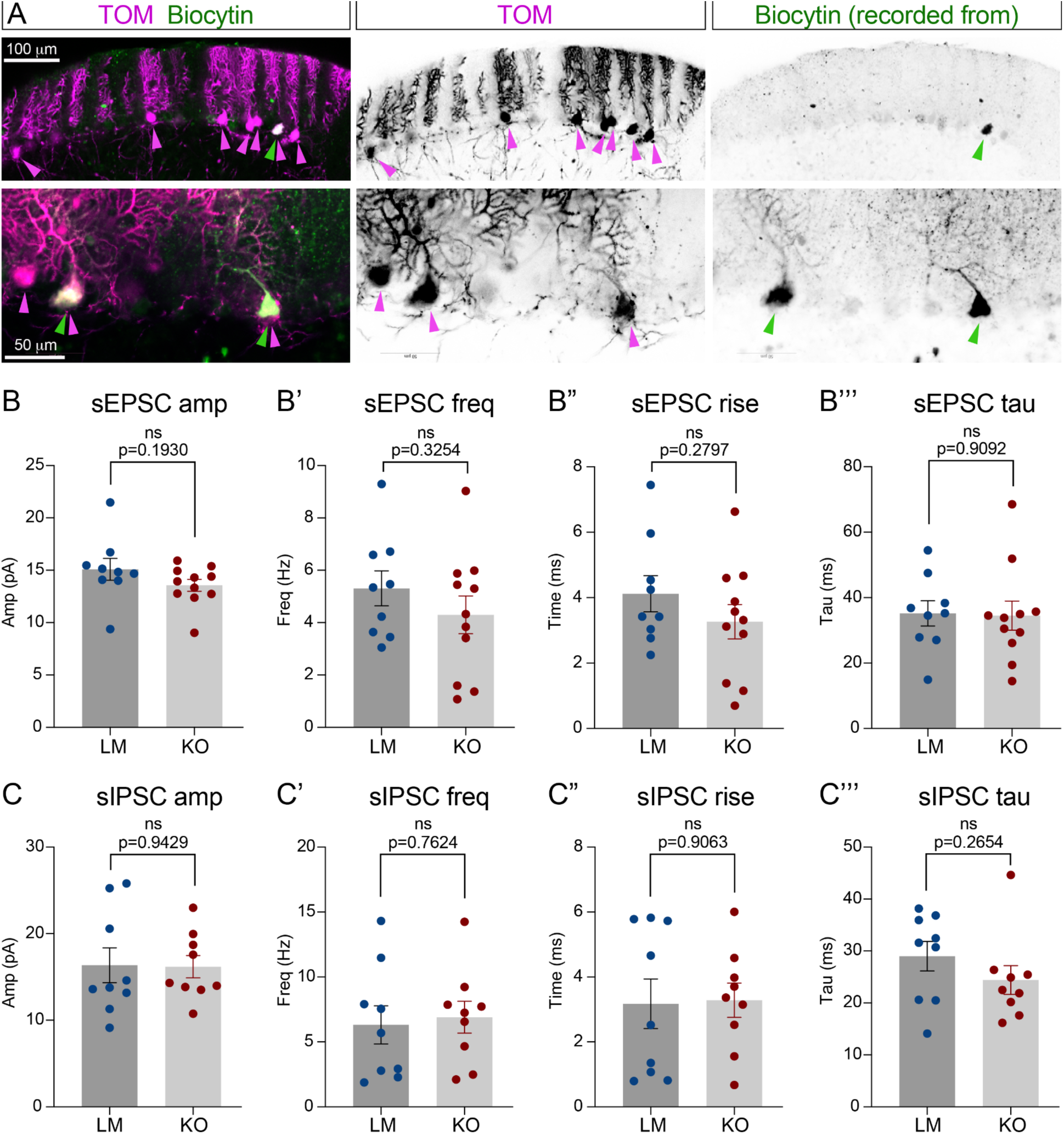
Spontaneous synaptic activity is normal in DAGLα-null Purkinje cells. (A) Representative images of PCs showing TOM (purple) and Biocytin (green) co-localization in recombined PCs from which the recordings were collected. Daglα KO PCs exhibit no differences in spontaneous excitatory postsynaptic currents (sEPSCs) parameters such as (B) amplitude, (B’) frequency, (B”) rise time, or (B””) tau. (C-C””) Spontaneous inhibitory postsynaptic currents (sIPSCs) are similarly unaffected. Columns show means (+-SEM). An unpaired student T-test was used to assess SEM and p-values.

**Supplemental Figure 4.**
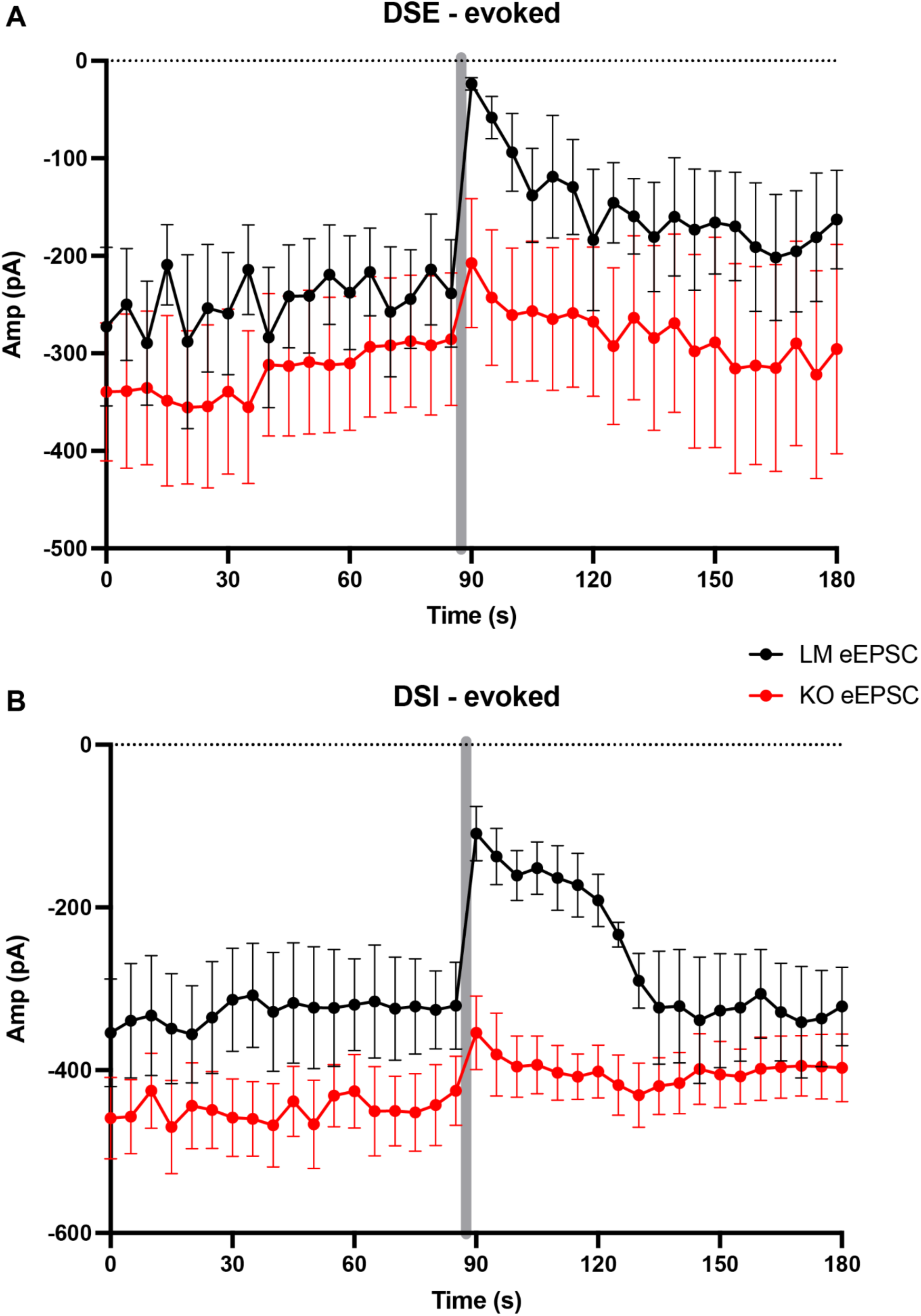
Evoked postsynaptic potentials before and after PC stimulation - not normalized. (A) eEPSC before and after PC stimulation in LMs and KOs. (B) eIPSC before and after PC stimulation in LMs and KOs.

